# Chemical tools to define and manipulate interferon-inducible Ubl protease USP18

**DOI:** 10.1101/2024.04.08.588544

**Authors:** Griffin J. Davis, Anthony O. Omole, Yejin Jung, Wioletta Rut, Ronald Holewinski, Kiall F. Suazo, Hong-Rae Kim, Mo Yang, Thorkell Andresson, Marcin Drag, Euna Yoo

## Abstract

Ubiquitin-specific protease 18 (USP18) is a multifunctional cysteine protease primarily responsible for deconjugating interferon-inducible ubiquitin-like (Ubl) modifier ISG15 from protein substrates. Here, we report the design and synthesis of activity-based probes (ABPs) capable of selectively detecting USP18 activity over other ISG15 cross-reactive deubiquitinases (DUBs) by incorporating unnatural amino acids into the C-terminal tail of ISG15. Combining with a ubiquitin-based DUB ABP, the selective USP18 ABP is employed in a chemoproteomic screening platform to identify and assess inhibitors of DUBs including USP18. We further demonstrate that USP18 ABPs can be utilized to profile differential activities of USP18 in lung cancer cell lines, providing a strategy that will help define the activity-related landscape of USP18 in different disease states and unravel important (de)ISGylation-dependent biological processes.

## Introduction

Deubiquitinases (DUBs), a family of proteases that cleave ubiquitin-like molecules (Ubls) from their substrate proteins, play critical roles in regulating protein turnover and non-degradative functions including DNA repair, protein complex formation, cellular trafficking, localization, and inflammation.^1^ DUBs are broadly implicated in many pathophysiological conditions including cancer, given their fundamental roles in various cellular processes.^2^

Among approximately 100 reported mammalian DUBs, ubiquitin-specific protease 18 (USP18) is primarily responsible for the cleavage of the interferon-inducible Ubl, interferon-stimulated gene 15 (ISG15) with exquisite specificity.^3^ In response to various cellular stresses, particularly viral infections and other immune stimuli, ISG15 conjugation (ISGylation) is mediated by the consecutive action of a three-step catalytic cascade in a similar manner to ubiquitylation,^4, 5^ and is counteracted by USP18 isopeptidase activity (Figure 1A). Although currently no consensus sequence for ISGylation has been defined, ISGylation of over 300 proteins has been reported in IFN-stimulated cells, including oncogenic and tumor suppressive proteins, as well as proteins involved in innate immune response, modulating their stability, activity, localization, and interactions.^6-8^ There is mounting evidence that ISGylation plays important roles in maintaining genome stability, cytoskeleton dynamics, autophagy, protein translation, and hypoxia/ischemic responses.^9^ While much attention has been given to the function of ubiquitin in protein degradation, it remains unclear how protein ISGylation is implicated in protein stability, degradation, or cross-talks with ubiquitylation. Independent of its deISGylating activity, USP18 also binds to IFN-α/β receptor 2 complex, where it competes with JAK1, thereby negatively regulating type I IFN signaling.^10, 11^ mRNA and protein levels of USP18 are found to be elevated in diverse cancers and USP18 loss has been shown to restrict tumor growth.^12, 13^ Despite these findings, the mechanistic contribution of this multifaceted protein to cancer development, progression, or survival remains largely understudied. USP18 can modulate the stability and functional activity of proteins important to tumor growth and antitumor immunity by regulating protein deISGylation and protein-protein interactions.^14-18^ As a major regulator of the IFN signaling network and T cell differentiation, USP18 can also affect inflammatory and immune responses within the tumor immune microenvironment.^19-21^ The effect of DUBs in cancer is often complex and highly context-dependent. Similarly, depending on the substrate landscape, subcellular localization, cell type, and physiologic states, each cancer type may respond differently to USP18 loss or inhibition. In addition, there are multiple mechanisms including protein binding and post-translational modifications (PTMs) that modulate DUB activity, the dysregulation of which has been observed in cancers.^22-24^ Recently, integrated multi-omics data revealed that the sensitivity of individual tumor lineages to different DUB knockouts is often not correlated with protein abundance,^25^ underscoring the importance of understanding the precise roles of DUBs for therapeutic intervention in cancer. There is a significant need for the development of chemical tools to understand the abundance, localization, substrate landscape, and activity of ISG15-specific protease USP18 in health and disease conditions, which will also provide insight into USP18-targeting therapeutic strategies.

**Figure 1.**
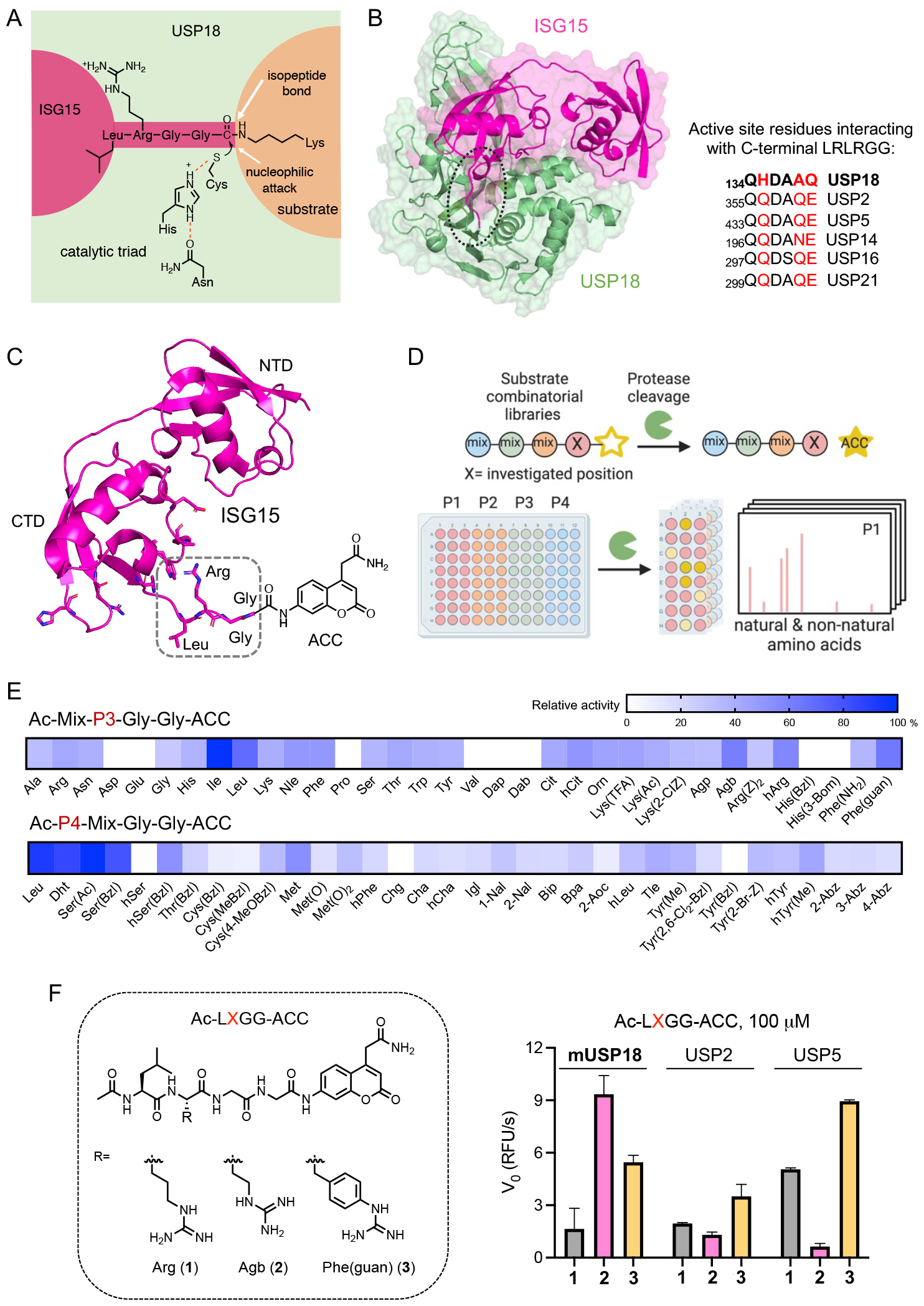
Fluorogenic substrates for USP18. (A) Catalytic triad of USP18. The isopeptide bond between the C-terminal Gly residue of ISG15 and the Lys residue of the substrate protein is hydrolyzed by the USP18 protease activity. (B) Structure of USP18 complexed with ISG15 (PDB: 5CHV). Active site residues of USP18 that interact with conserved C-terminal LRLRGG of ISG15 are highlighted in comparison to other ISG15 reactive DUBs. (C) Structure of ISG15-based ACC-labeled substrate. The C-terminal LRGG motif is highlighted. (D) Schematic overview of HyCoSuL screening. (E) Substrate profiles presented as heatmaps indicating the percentage of relative cleavage measured by a fluorescent signal produced upon USP18 binding with two sub-libraries, Ac-Mix-P3-Gly-Gly-ACC and Ac-P4-Mix-Gly-Gly-ACC (where Mix represents an equimolar mixture of natural amino acids). (F) Structures (left panel) and rate of hydrolysis (right panel) of selected tetrapeptide fluorogenic substrates. mUSP18 (10 μM), USP2 (2 μM), and USP5 (1 μM) were incubated with 100 μM of Ac-LXGG-ACC substrates for 1 h at RT and initial release of fluorescent ACC (V_0_, RFU/s) by enzyme was measured at Ex: 360 nm / Em: 460 nm. Agb: 2-amino-4-guanidino-butyric acid, hArg: homoarginine, Phe(guan): guanidino-phenylalanine.

A chemoproteomic method termed activity-based protein profiling (ABPP) that uses activity-based probes (ABPs) is a powerful platform to map proteome-wide reactive proteins and evaluate the functional states of enzymes in complex biological systems.^26, 27^ Using ABPs that mimic native ubiquitin but are modified with an electrophilic warhead at the C-terminus to label the active site cysteines irreversibly, ABPP has led to the identification of new DUB families and characterization of DUB activity across conditions.^28^ In a recent study, an ISG15-based ABP was used to detect deISGylating enzymes in human cell lysates and identified USP16 as an ISG15 cross-reactive DUB in addition to the previously-characterized DUBs such as USP5 and USP14.^29-32^ Competitive ABPP has also been implemented to screen DUB inhibitors.^33, 34^

Here, we report the design and synthesis of an ABP that selectively detects deISGylase activity of USP18. Through screening of a hybrid combinatorial substrate library (HyCoSuL),^35^ we identified mutations in the LRGG tail of ISG15 that enhance selective binding to USP18 over other ISG15 cross-reactive DUBs. We incorporated these mutations into a C-terminal domain of murine ISG15 using a total chemical synthesis approach and evaluated their efficiency and selectivity in cell lysate conditions. These selective USP18 ABPs were employed to identify WP1130 as an inhibitor of USP18 in a workflow that can be adapted as a chemoproteomic screening platform for discovering and characterizing DUB inhibitors. We further utilized them for profiling USP18 activity in cell lines derived from lung carcinoma. Our study indicates that the differential activity of USP18 can be measured using this approach to study its activity-related biological functions in cancer.

## Results and Discussion

### Design and synthesis of mISG15_CTD_-based probes for selective detection of USP18

In general, a DUB ABP consists of three components: a ubiquitin-like protein (recognition element), an electrophile such as vinyl sulfone (VS) or propargyl (PA) group (reactive warhead), and a fluorophore/affinity handle (reporter tag).^28^ Upon Ubl binding to DUB, the reactive electrophile is covalently attached to the active site cysteine in an enzyme-catalyzed reaction and the probe labeling reports DUB activity. To develop an ABP for USP18, we first explored requirement for the recognition element of the ISG15 sequence. ISG15 consists of two Ubl domains connected by a short hinge region and contains the canonical LRGG motif at its C-terminus, which is required for conjugation to its targets.^36^ When we tested Ub, ISG15 (ISG15_FL_), and the C-terminal Ubl domain of ISG15 (ISG15_CTD_) in vitro, we observed that USP18 prefers and efficiently binds to ISG15 compared to Ub as previously reported, and ISG15_CTD_ is sufficient for binding to USP18 (Figure S1A).^37^ We also found that mouse ISG15 (64% sequence identity to human ISG15) binds well to human USP18. When native ISG15 is used as a recognition element, however, these ISG15-based probes have been reported to label not only USP18 but also other ISG15 cross-reactive DUBs such as USP2, USP5, USP14, and USP16.^30, 31^ We found that, indeed, USP5 was labeled by both hISG15_FL_ and mISG15_CTD_-based probes (Figure S1B).

To develop an ABP that specifically reports on the deISGylating activity of USP18, we hypothesized that creating mutations in certain regions of ISG15 while keeping key residues that confer specificity over Ub,^37^ may optimize intermolecular contacts and direct it toward USP18 with enhanced binding affinity and selectivity. To assess whether mutations in the conserved C-terminal LRGG motif can be tolerated or even enhance interaction with USP18 based on the subtle differences in the active site residues of USP18 compared to other ISG15 cross-reactive DUBs (Figure 1B), we performed positional scanning of a hybrid combinatorial substrate library (HyCoSuL),^35, 38^ which allows rapid substrate profiling for amino acid preferences including natural and unnatural amino acids at the sites adjacent to the scissile bond (Figure 1C-D). P3 and P4 sub-libraries of fluorogenic tetrapeptide substrates that bear Gly residues at the P1 and P2 and a 7-amino-4-carbamoylmethylcoumarin (ACC) fluorophore at the P1’ position were screened with USP18. We found that USP18 recognizes and cleaves several tetrapeptide substrates that contain unnatural amino acids, for example, Agb, hArg, Phe(guan) at the P3, and Dht, Ser(Ac), Ser(Bzl) at the P4 position (Figure 1E). We then synthesized seven individual fluorogenic substrates (i.e., Ac-LXGG-ACC and Ac-XRGG-ACC) and measured the rate of hydrolysis (Figure S2). The Ac-Leu-Agb-Gly-Gly-ACC substrate (**2**) was cleaved faster than Ac-LRGG-ACC (**1**) by mUSP18 but showed decreased rates of cleavage by USP2 and USP5, while Ac-Leu-Phe(guan)-Gly-Gly-ACC substrate (**3**) was efficiently cleaved by all three enzymes (Figure 1F). This suggests that USP18 can recognize amino acids other than the canonical Leu-Arg in the P4-P3 positions while replacing the P3 Arg with Agb might enhance selectivity towards USP18.

Since tetrapeptide sequences are not sufficient to fully engage USP18, we synthesized the C-terminal domain of mISG15 using standard solid-phase peptide synthesis to incorporate unnatural amino acids, warheads, and reporter tags in a straightforward manner with chemical flexibility (Figure 2A). Briefly, we used Fmoc-Gly TentaGel Trt resin and sequentially deprotected Fmoc and coupled the following amino acids in the presence of PyBOP and DIPEA in NMP.^39^ To improve stability and avoid dimerization, Cys144 was replaced by Ser. Mutant probes were generated by substituting Arg153 with Agb, hArg, or Phe(guan) accordingly. Once mISG15_CTD_ (Leu80∼Gly154) was assembled, the N-terminal end was either acetylated or conjugated with Cy5 fluorophore for visualization or with biotin for affinity purification. After cleavage from the solid support, the C-terminal end was further functionalized with a propargylamine warhead, replacing Gly155, for covalent conjugation.^40^ The identity and purity of each probe were confirmed by LC-MS analysis (Figure 2B). Our SPR analysis of Ac-mISG15_CTD_-OH indicated that both Agb and Phe(guan) mutants retain binding affinity toward USP18 (Figure S3).

**Figure 2.**
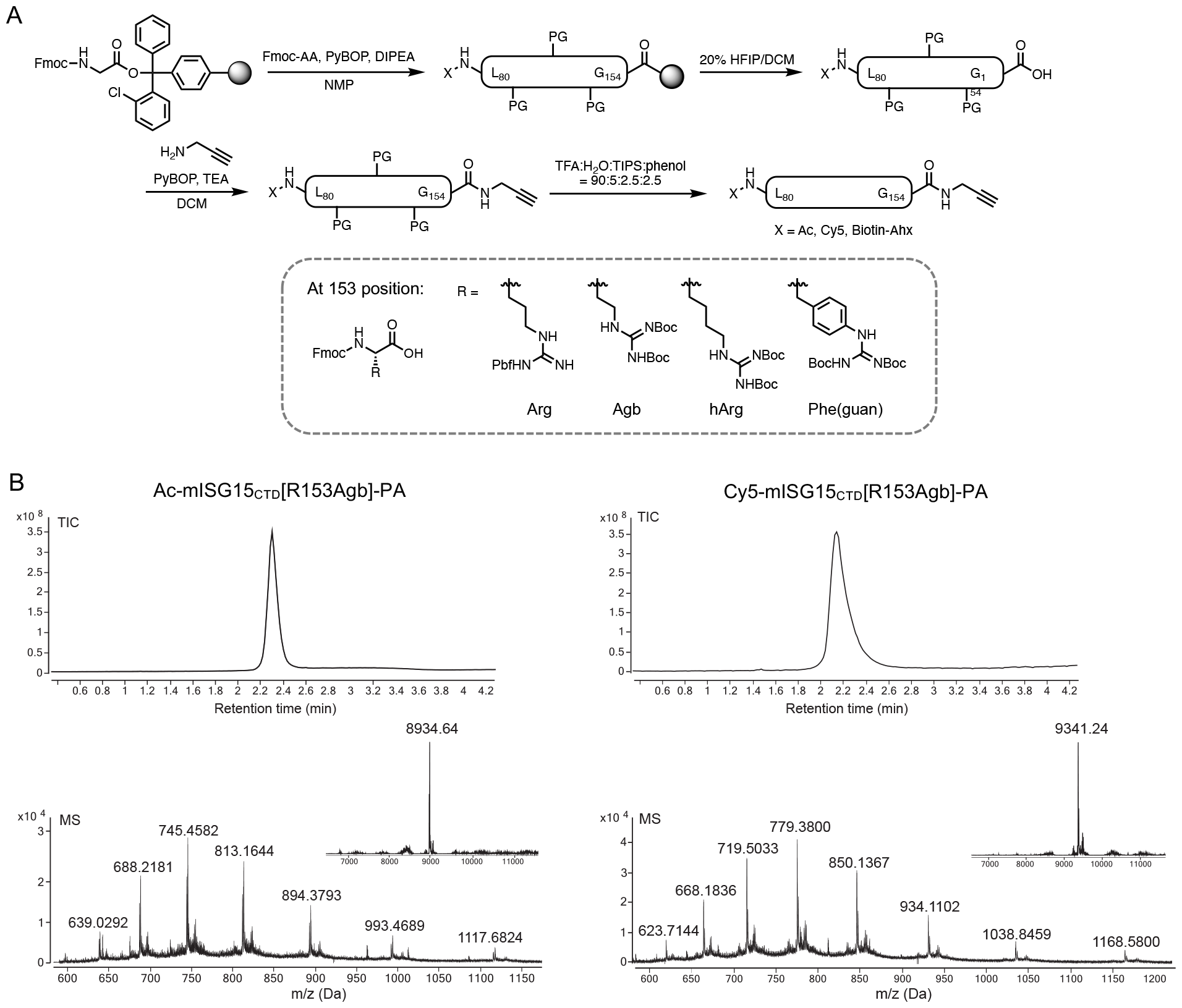
Preparation of mISG15_CTD_-based probes. (A) Synthesis of C-terminal domain of mouse ISG15 and ISG15 variants. Leu80∼Gly154 sequence was assembled using SPPS and the N-terminal end was modified by either an acetyl group, fluorescent cyanine dye, or biotin affinity handle. After the cleavage from the solid support, the C-terminal end was functionalized with propargylamine (PA) replacing Gly155. Arg153 was varied with unnatural amino acids such as Agb (2-amino-4-guanidino-butyric acid), hArg (homoarginine), Phe(guan)(guanidino-phenylalanine). (B) LC-MS analysis of synthetic Ac-mISG15_CTD_-PA and Cy5-mISG15_CTD_-PA probes.

### Reactivity and selectivity assessment of mISG15_CTD_-based probes

We then performed gel-based assays to assess the reactivity and selectivity of mutant ISG15 probes. When incubated with both mouse and human recombinant USP18, all tested Ac-mISG15_CTD_-PA probes showed efficient labeling of the active enzyme, judging from the appearance of a distinct, higher molecular weight protein band owing to covalent bond formation (Figure 3A). Importantly, the Agb and Phe(guan) mutant probes showed diminished to marginal interaction with USP5, while the wild type (WT) and hArg mutant displayed significant conjugation (Figure 3B), revealing that subtle change in the tail of ISG15, particularly at the 153 position (varying the length of the side chain by no more than two methylene carbons), has significant consequences for its molecular interaction.

**Figure 3.**
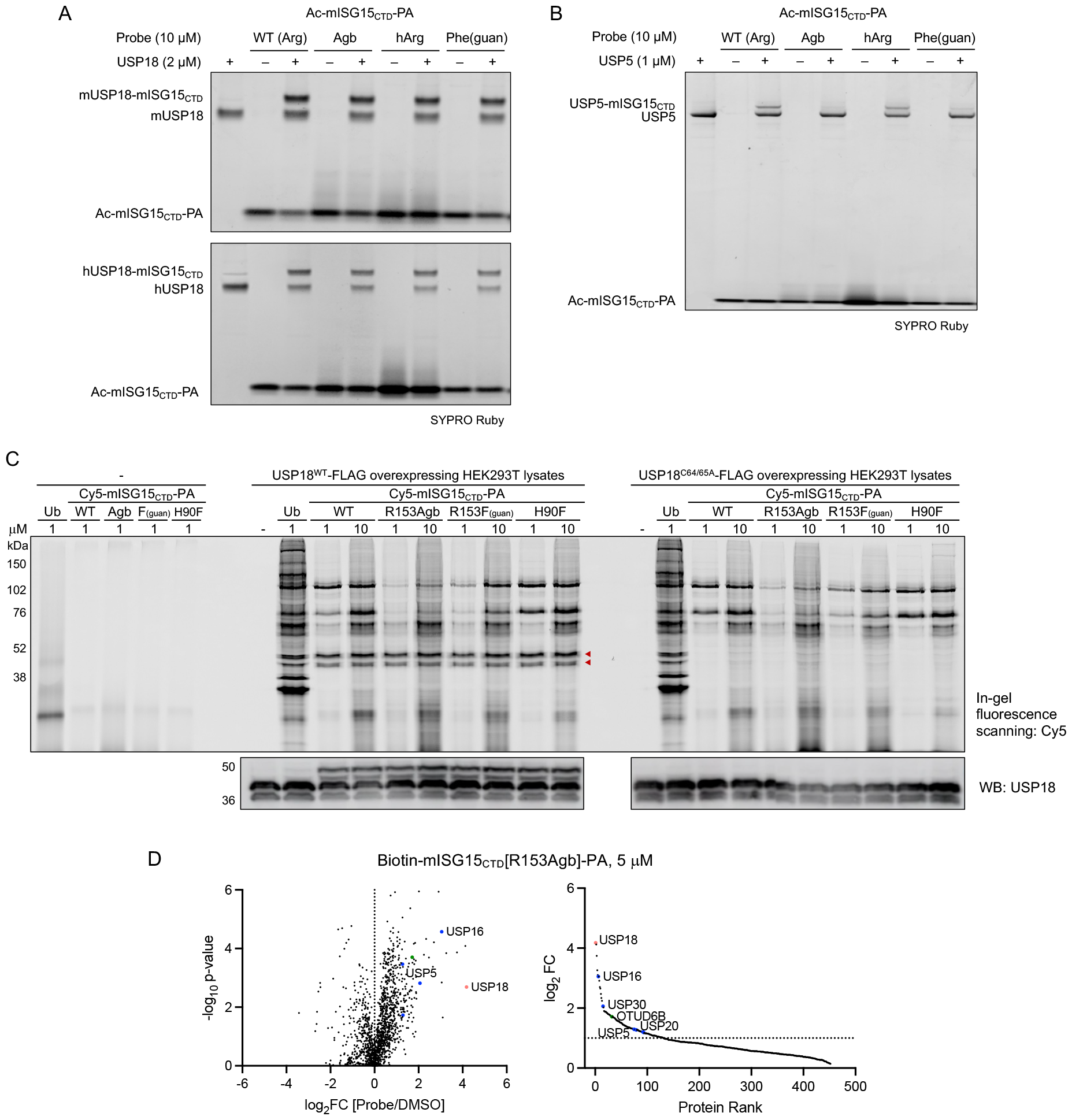
Reactivity and selectivity of mISG15_CTD_-based probes. (A-B) Recombinant mouse and human USP18 (2 μM) or USP5 (1 μM) were incubated with Ac-mISG15_CTD_-PA probes (10 μM) for 3 h at RT. Protein samples were analyzed by SDS-PAGE and SYPRO Ruby staining. (C) WT or C64/65A mutant of USP18-FLAG overexpressing HEK293T cell lysates were incubated with Cy5-mISG15_CTD_-PA probes at indicated concentrations for 3 h at RT. Protein samples were analyzed by SDS-PAGE and in-gel fluorescence scanning for Cy5 signal. Red marks indicate the appearance of new protein bands only in USP18^WT^ overexpressing cell lysates after labeling by probes corresponding to the expected molecular weight of USP18−probe conjugate. Expression of USP18 was confirmed by western blotting. (D) Volcano (left) and protein rank (right) plots of quantitative proteomic analysis of streptavidin beads pulldowns after labeling of USP18-FLAG overexpressing HEK293T cell lysates by Biotin-mISG15_CTD_[R153Agb]-PA probe (5 μM, 3 h, RT) showing significantly enriched proteins (log_2_ ratio > 1, p-value ≤ 0.05). USP18 is marked and other DUBs are colored based on subfamilies.

To visualize DUB–probe conjugates in complex proteomes, we first used Cy5-Ubl-PA probes in lysates from HEK293T cells (Figure S4). We included an H90F mutant of mISG15_CTD_ (a mutation of mISG15_CTD_ not in the tail region) that was computationally predicted to increase binding to USP18 through hydrophobic interaction. The Ub probe demonstrated clear labeling of many proteins, while the WT mISG15_CTD_ probe showed comparatively less but still strong labeling of several proteins as expected. With no detectable basal expression of USP18, Arg153Agb and Arg153Phe(guan) mutant probes showed decreased background labeling relative to the WT, whereas the H90F mutant probe showed increased background labeling as noted by a more intense band with molecular weight near 76 kDa. In the USP18^WT^ overexpressing HEK293T cell lysates, two additional protein bands appeared, corresponding to covalent adducts of two USP18 isoforms^41^ (red arrows in Figure 3C), which was elaborated by western blot analysis. Satisfyingly, when the labeling assay was performed with the lysates of HEK293T cells overexpressing catalytically inactive USP18^C64A/C65A^ (a double mutation of catalytic (C64) and adjacent (C65) cysteines),^16^ there was no USP18–probe conjugate detected, confirming that apparent labeling of USP18 by ABPs is through covalent conjugation in a catalytic cysteine-dependent manner. Overall, the Arg153Agb mutant appeared to be slightly more selective for USP18 than the Arg153Phe(guan) mutant, as concluded from less intense labeling of other ISG15 cross-reactive DUBs. From preincubation of USP18^WT^ overexpressing HEK293T cell lysates with either the WT or Arg153Agb mutant of Ac-mISG15_CTD_-PA prior to labeling with Cy5-mISG15_CTD_[WT]-PA, we observed that the R153Agb mutant selectively blocked the USP18 labeling while the WT probe competed for other proteins in addition to USP18 (Figure S5), again confirming enhanced selectivity of Arg153Agb for USP18 compared to the WT.

For parallel affinity pulldown and LC-MS/MS analysis, we also devised biotin ABPs. The biotin probes were evaluated with recombinant enzymes first to ensure that N-terminal modification with biotin retains the same binding and selectivity behavior. The biotin-mISG15_CTD_[R153Agb]-PA probe showed sufficient binding to USP18 (Figure S6A) with minimal binding to USP5 (Figure S6B). Immunoblotting analysis of pulldown experiments in USP18 overexpressing HEK293T cell lysates confirmed the enrichment of active USP18 by the R153Agb mutant probe (Figure S6C). The probe-labeled, streptavidin-enriched proteins were then subjected to on-bead digestion, isobaric tandem mass tag (TMT) labeling, and LC-MS/MS analysis (Figure S7A). From HEK293T cell lysates, the biotin-Ub-PA probe was found to enrich 36 active DUBs compared to the DMSO control (log_2_ fold change > 1, p-value ≤ 0.05 in triplicates) (Figure S7B and Table S1). Of the enriched DUBs, 24 belong to the USP family. The biotin-mISG15_CTD_[WT]-PA probe enriched far fewer proteins, but 3 members of the USP family − USP5, USP14, and USP16 − were highly enriched. This finding is consistent with the previous identification of these DUBs to cross-react with ISG15 in vitro.^29^ In contrast, the R153Agb probe showed significantly reduced enrichment of USP binding partners when there was no detectable USP18 present. We then repeated the ABPP experiment with USP18^WT^-FLAG overexpressing HEK293T cell lysates. Considering the lower abundance of USP18 compared to other DUBs, we opted to treat with a relatively high probe concentration (5 μM) to ensure proteome coverage. As expected, USP18 was the protein most highly enriched by the R153Agb probe followed by USP16 (Figure 3D). Very minimal or no significant enrichment was observed for USP5 and USP14, respectively. Overall, the ABPP data recapitulate the selectivity profile of our USP18 ABP and establish its utility in quantitative chemoproteomics as a chemical tool to selectively report USP18 activity in various cell lines.

### USP18 inhibitor screening using a chemoproteomic platform

With this selective USP18 ABP, we sought to perform competitive ABPP to evaluate target engagement and selectivity of DUB inhibitors (Figure 4A)^33, 34, 42^ Since USP18 does not react with Ub, USP18 is excluded in DUB inhibitor screening when using Ub-based probes despite its biological importance and potential as a drug target. However, we found that several reported DUB inhibitors, including WP1130 (a known inhibitor of USP5, USP9X, USP14, USP24, and UCH37, Figure 4B),^43-45^ inhibit USP18 protease activity in an ISG15-Rho assay (Figure S8). We first validated that WP1130 binding prevents the labeling of recombinant USP18 by the WT ISG15-based covalent probe (Figure 4C). When USP18 overexpressing HEK293T cell lysates were preincubated with WP1130 for 1 h and treated with each Cy5-Ubl probe (1 μM) for 2 h, we observed a dose-dependent decrease in fluorescent intensity of conjugated USP18 and some other DUBs with ISG15-based probes and the Ub-based probe, respectively (Figure 4D). Similarly, when a cocktail of biotin probes (1 μM of Biotin-Ub-PA and 5 μM of Biotin-mISG15_CTD_[R153Agb]-PA) was used to label active DUBs including USP18, immunoblotting analysis of samples preincubated with WP1130 showed dose-dependent inhibition of USP18 activity (Figure 4E). Quantitative MS analysis also corroborated western blotting results with a surprisingly narrow reactivity scope. Among 48 active DUBs detected using a cocktail of probes (log_2_FC > 1, p-value ≤ 0.05 in triplicates, Figure 4F and Table S2), WP1130 binding was found to compete for probe labeling of 4 DUBs – USP22, USP42, USP33, and USP18 – of which USP18 was the most efficiently blocked (Figure 4G and S9). Although not specific, we thus report WP1130 as a promising, first example of small-molecule USP18 inhibitors with the potential for optimization. The reason we were unable to detect other DUBs that WP1130 is known to target might be due to the use of USP18 overexpressing lysates (as we observed WP1130 competitively blocks the labeling of USP9X and USP24 in HEK293T cell lysates, data not shown) and the reversible nature of modification by WP1130. Nonetheless, our study demonstrates that utilizing this USP18 ABP in conjunction with a Ub-based DUB probe in a chemoproteomic screening platform holds promise as a tractable workflow for global, competitive profiling of DUB inhibitors to perform simultaneous hit identification and selectivity assessment. Although we used the USP18 overexpressing system as a model, this screening approach can be applied to native or disease models of interest.

**Figure 4.**
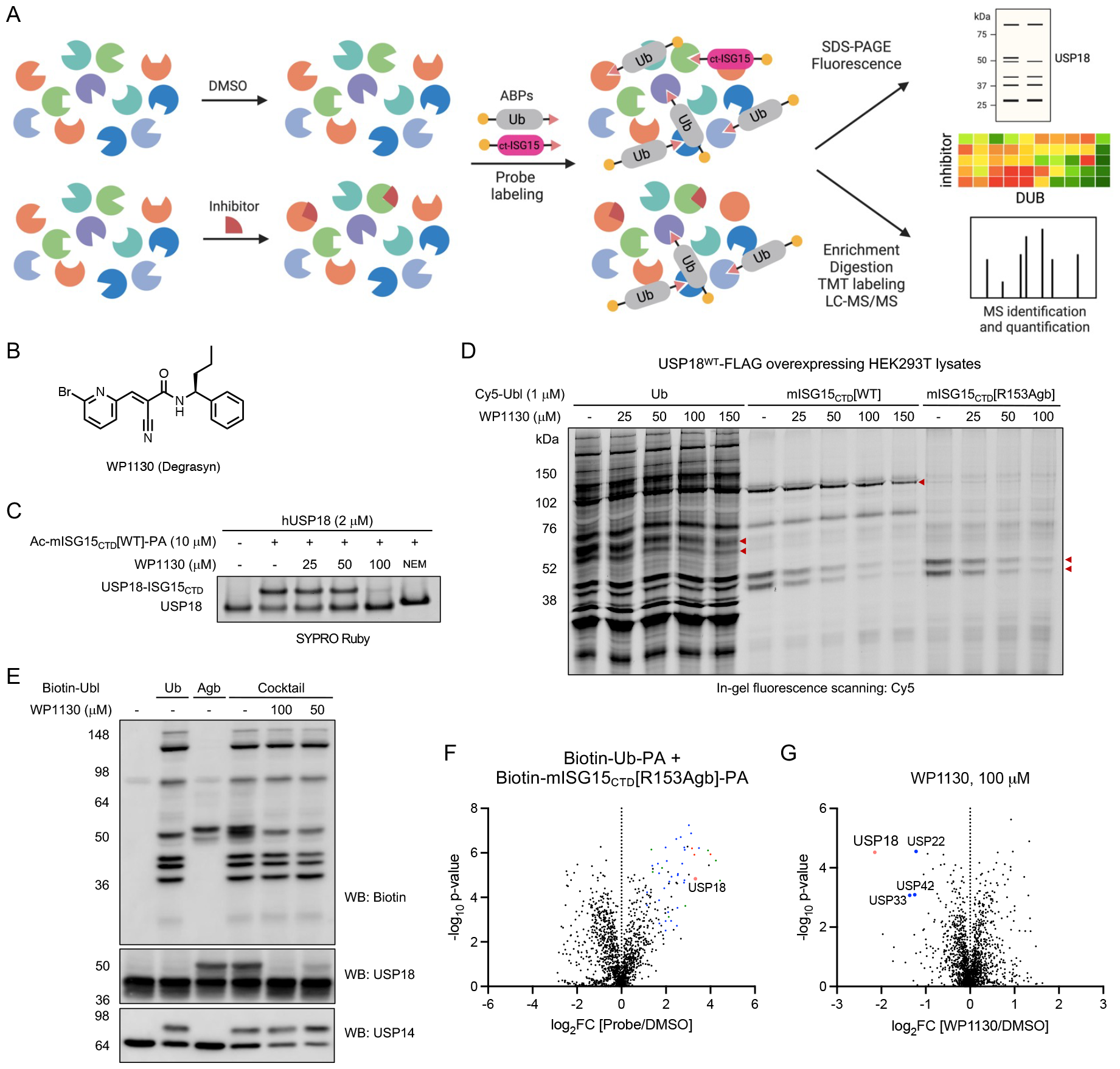
USP18 inhibitor assay by competitive activity-based protein profiling. (A) Schematic workflow for quantitative activity-based protein profiling. Lysates were treated with DMSO or compound, followed by treatment with an activity-based DUB probe. Labeled proteins were enriched using biotin-streptavidin pull-down, digested, and analyzed by LC-MS/MS. (B) Chemical structure of WP1130. (C) Recombinant human USP18 (2 μM) was pretreated with WP1130 for 30 min and incubated with Ac-mISG15_CTD_[WT]-PA probe (10 μM) for 2 h at RT. Protein samples were analyzed by SDS-PAGE and SYPRO Ruby staining. (D) USP18^WT^-FLAG overexpressing HEK293T cell lysates were preincubated with WP1130 for 1 h followed by labeling with 1 μM of Cy5-Ubl-PA for 2 h at RT. Protein samples were analyzed by SDS-PAGE and in-gel fluorescence scanning for Cy5 signal. (E-G) USP18^WT^-FLAG overexpressing HEK293T cell lysates were preincubated with WP1130 for 2 h followed by labeling with 1 μM of Biotin-Ub-PA or 5 μM of Biotin-mISG15_CTD_[R153Agb]-PA or a cocktail of probes (1 μM of Biotin-Ub-PA + 5 μM of Biotin-mISG15_CTD_[R153Agb]-PA) for 3 h at RT. Protein samples were analyzed by SDS-PAGE and immunoblotting (E). Volcano plots of quantitative proteomic analysis of streptavidin beads pulldowns after pretreating cell lysates with either DMSO (F) or 100 μM of WP1130 (G) and labeling by a cocktail of probes. Significantly enriched proteins are shown (log_2_ ratio > 1, p-value ≤ 0.05). USP18 is marked and other DUBs are colored based on subfamilies.

### Activity-based profiling of USP18 in lung cancer cell lines

As activity-based protein profiling can determine DUB activities in different pathophysiological conditions,^46, 47^ we chose to profile USP18 activity in lung carcinoma cell lines based on the analysis of the expression and gene effect of USP18 observed from the Broad DepMap database (Figure S10). A small number of cell lines, A549, H358, H2030, and H1650 lung cancer lines were selected and first evaluated for their USP18 expression. We found that these cell lines express high levels of both full-length and truncated USP18 upon IFN-β stimulation (Figure 5A). Under basal conditions, H358 and H1650 cells appeared to have higher levels of USP18 expressed compared with A549 cells. Cell lysates were then incubated with Cy5-mISG15_CTD_-PA probe (1 μM, 3 h) and analyzed by SDS-PAGE and in-gel fluorescence scanning (Figures 5B and S11). We observed fluorescently labeled protein bands around 40∼50 kDa in IFN stimulation conditions across cell lines, indicative of USP18−probe conjugates with increased intensity compared to non-IFN stimulation conditions. The fluorescent intensity of the USP18sf (truncated isoform)−probe conjugate band measured by densitometry indicated a slightly higher activity of USP18 in H2030 cells. This approach of measuring the differential activity of USP18 combined with expression analysis can help investigate cancer types and states in which USP18 is highly activated and provide insights into posttranslational regulation of USP18 and the cellular context in which USP18 inhibitors can have translational potential.

**Figure 5.**
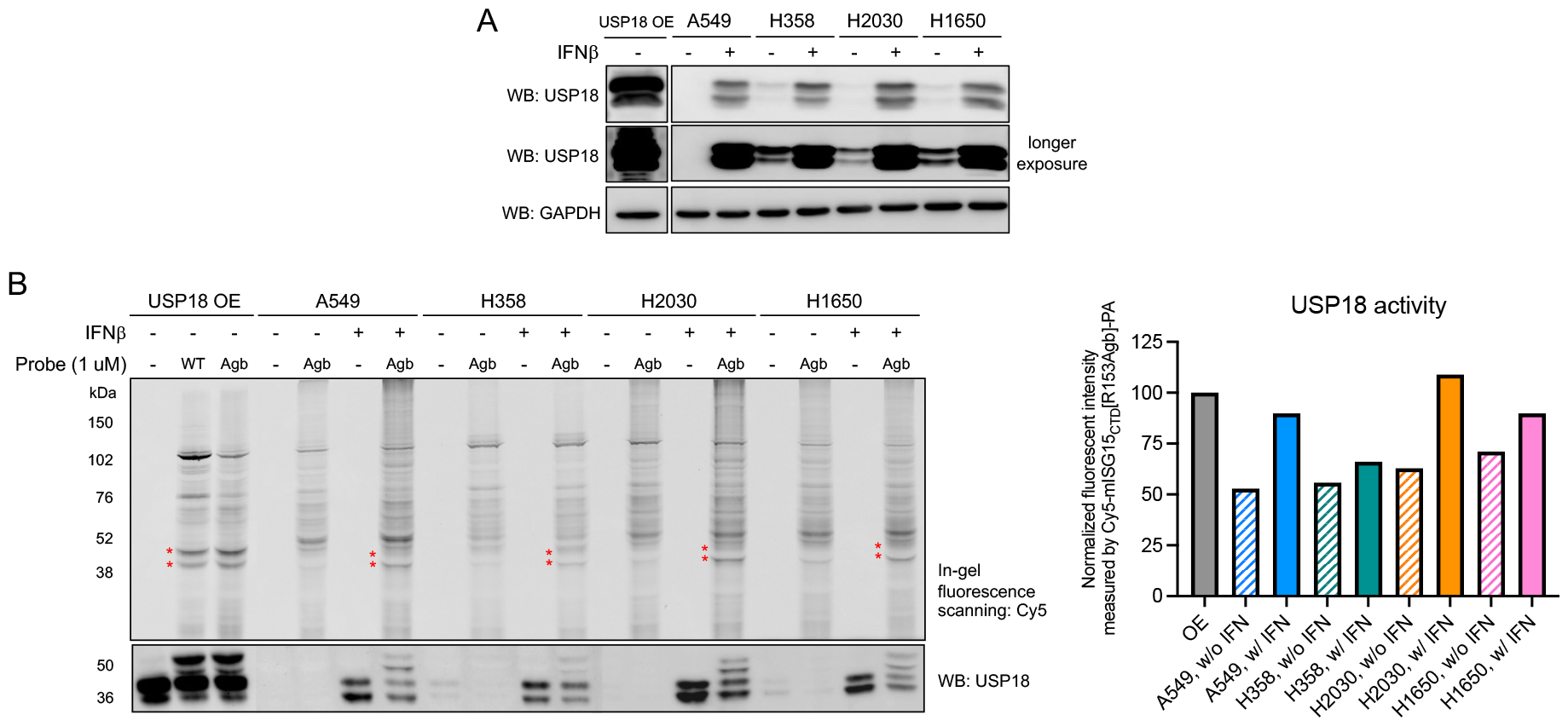
USP18 expression and activity profiles in lung cancer cell lines. (A) Constitutive and induced USP18 expression levels upon IFN-β stimulation (48 h) across lung cancer cell lines. (B) Each cell lysates were incubated with 1 μM of Cy5-mISG15_CTD_[R153Agb]-PA probe for 3 h at RT. Left, protein samples were analyzed by SDS-PAGE and in-gel fluorescence scanning for Cy5 signal. Red marks indicate the appearance of USP18−probe conjugates. Expression of USP18 was confirmed by western blotting. Right, normalized intensity of fluorescent protein band corresponding to the USP18sf−probe conjugate.

## Conclusion

Despite significant advancement in developing DUB probes, selective chemical tools to study a single DUB are limited. In an effort to develop chemical tools selectively detecting deISGylating activity of USP18 that would allow for the investigation of important (de)ISGylation-dependent biological processes, we identified non-canonical amino acids that can be incorporated into mISG15_CTD_ to preserve reactivity with USP18 while improving selectivity over other ISG15 cross-reactive DUBs such as USP5 and USP14. To understand this gained selectivity, we compared structures of DUB-Ubl complexes within the protease active site. In crystal structures of Ub complexed with USP5 (Figure S12B) and USP14 (Figure S12C), the C-terminal Ub LRGG motif is arranged such that the side chain of R74 forms numerous hydrogen bonds with backbone and side chains of surrounding active site residues (e.g., E427, Q433, Q434 in USP5 and Q196, Q197 in USP14). Shortening the side chain of arginine by a single methylene carbon would disrupt these interactions. On the other hand, unlike Q434 of USP5 or Q197 of USP14, H135 of USP18 is oriented away from the Ubl tail and the side chain of R153 participates in a hydrogen bond interaction in a way that would not be disrupted by substitution with Agb (Figure S12A). Solving the structure of USP18 bound with our Agb mutant probe would provide insight into how to engage major interactions within the active site and determine ISG15 mutants that are susceptible to hydrolysis only by USP18 when conjugated to the target substrate protein.

Using an ABPP platform, our work supports the recent discovery of USP16 as an ISG15 cross-reactive DUB^29, 30^ and demonstrates the ability to detect USP18 activity in different cell conditions. Moving forward, this strategy should allow us to profile USP18 activities in various cellular contexts and disease states, and eventually elucidate the contribution of its deISGylating activity to the disease phenotype. Combined with the Ub-based DUB activity profiling data, the ISG15-based activity profiling of deISGylases and USP18 will help comprehensively investigate DUBs and Ubl proteases, define the activity-related landscape of Ubl proteases in different disease states, and unravel important Ub and Ubl-dependent biological processes. In addition, adaptable workflow and application in inhibitor screening using this USP18 will probe provide a road map for developing DUB and USP18 inhibitors.

## Associated Content

### Supporting Information

Additional figures, detailed procedures for all experiments and computations, specifics for reagents and instruments used, and synthetic details (PDF) Proteomics data (Tables S1-2) (XLSX)

## Data availability

TMT-MS data have been deposited at MassIVE with accession number MSV000094406 (ftp://MSV000094406@massive.ucsd.edu).

## Acknowledgments

The cloning, expression, and purification of mouse and human USP18 were done at the Protein Expression Laboratory, Frederick National Laboratory for Cancer Research, and the authors are grateful to Dr. Jane Jones’s team for their assistance. The authors thank Dr. Ji Luo’s group (Laboratory of Cancer Biology and Genetics, CCR, NCI) for providing two lung cancer cell lines used in this study (H358 and H2030). This work was supported by the Intramural Research Program of the NIH, National Cancer Institute, Center for Cancer Research (ZIABC011963), National Science Center grant 2015/17/N/ST5/03072 (Preludium 9) in Poland (W.R.) and the “TEAM/2017-4/32” project, which is carried out within the TEAM program of the Foundation for Polish Science, cofinanced by the European Union under the European Regional Development Fund (M.D.). The authors gratefully thank the staff members at the CCR-Frederick Biophysics Resource (BR) for their assistance, technical consultation, and instrument maintenance.

---

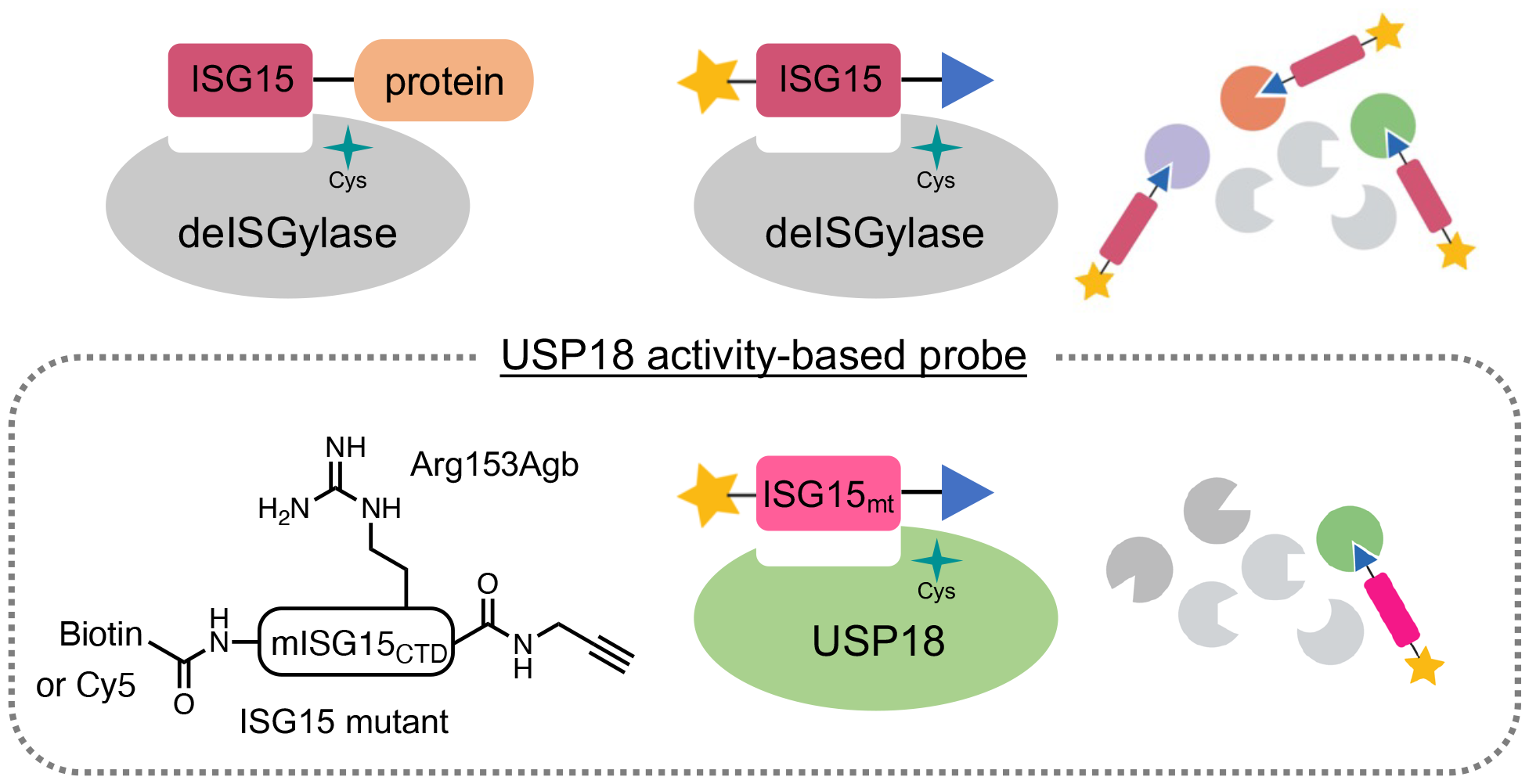

