## Supplemental information for "Chemical tools to define and manipulate interferon-inducible Ubl protease USP18"

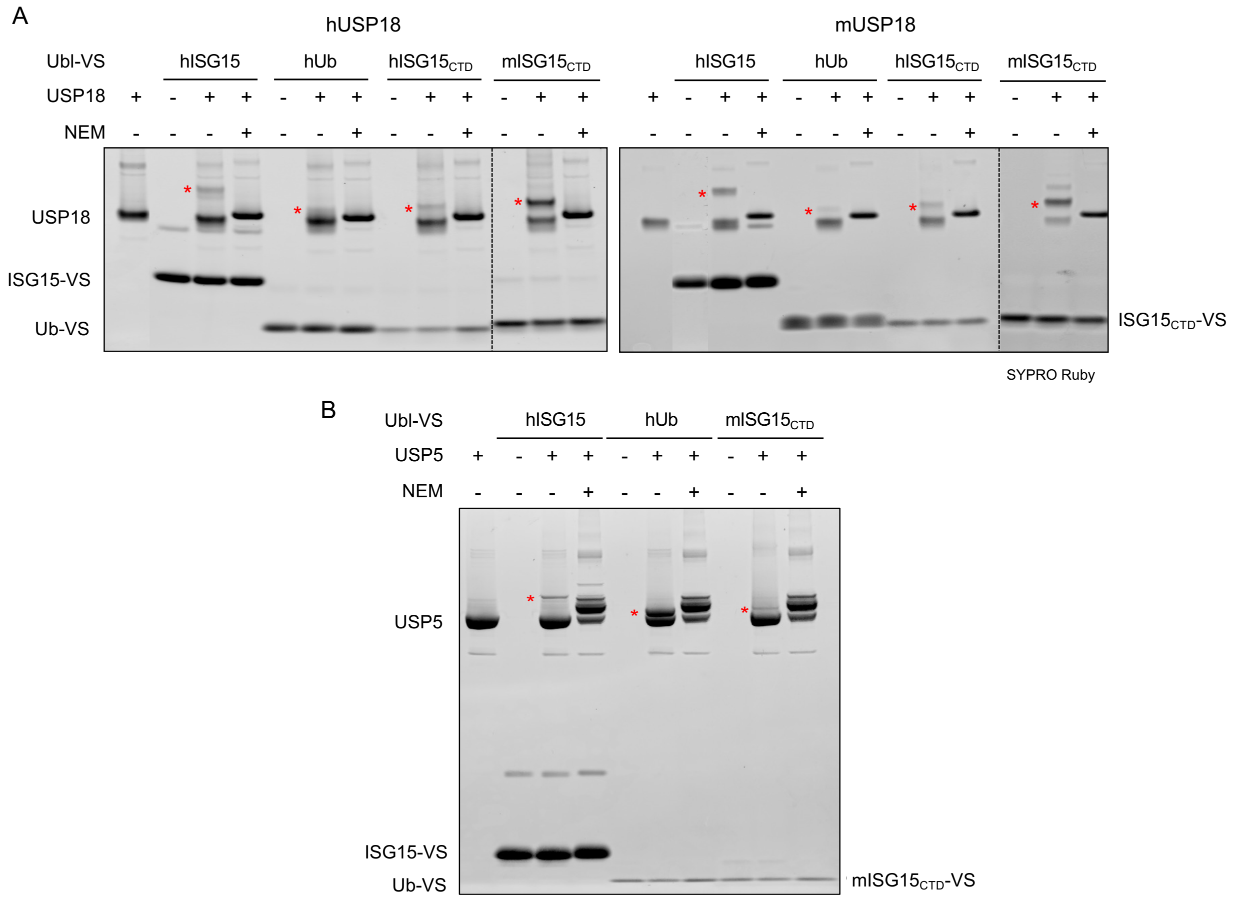


Figure S1. Reactivity Ub-VS and ISG15-VS towards USP18 and USP5. (A) Recombinant human and mouse USP18 (2 μM) was incubated with hISG15-VS (5 μM), hUb-VS (20 μM), h/mISG15$_{CTD}$-VS (10 μM) for 3 h at RT. (B) Recombinant human USP5 (1.5 μM) was incubated with each Ubl-VS (5 μM) for 3 h at RT. NEM pretreatment (10 μM for 10 min at RT) was performed where indicated. Protein samples were analyzed by SDS-PAGE and SYPRO Ruby staining.

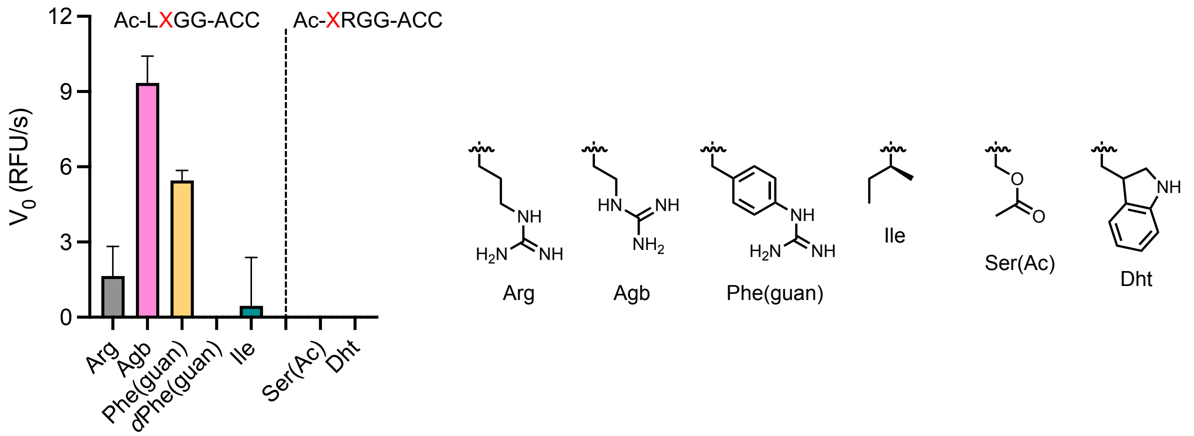


Figure S2. Rate of hydrolysis of selected tetrapeptide fluorogenic substrates. mUSP18 (10 μM) was incubated with 100 μM of substrates for 1 h at RT and the initial release of fluorescent ACC ($V_0$, RFU/s) by enzyme was measured at Ex: 360 nm / Em: 460 nm.

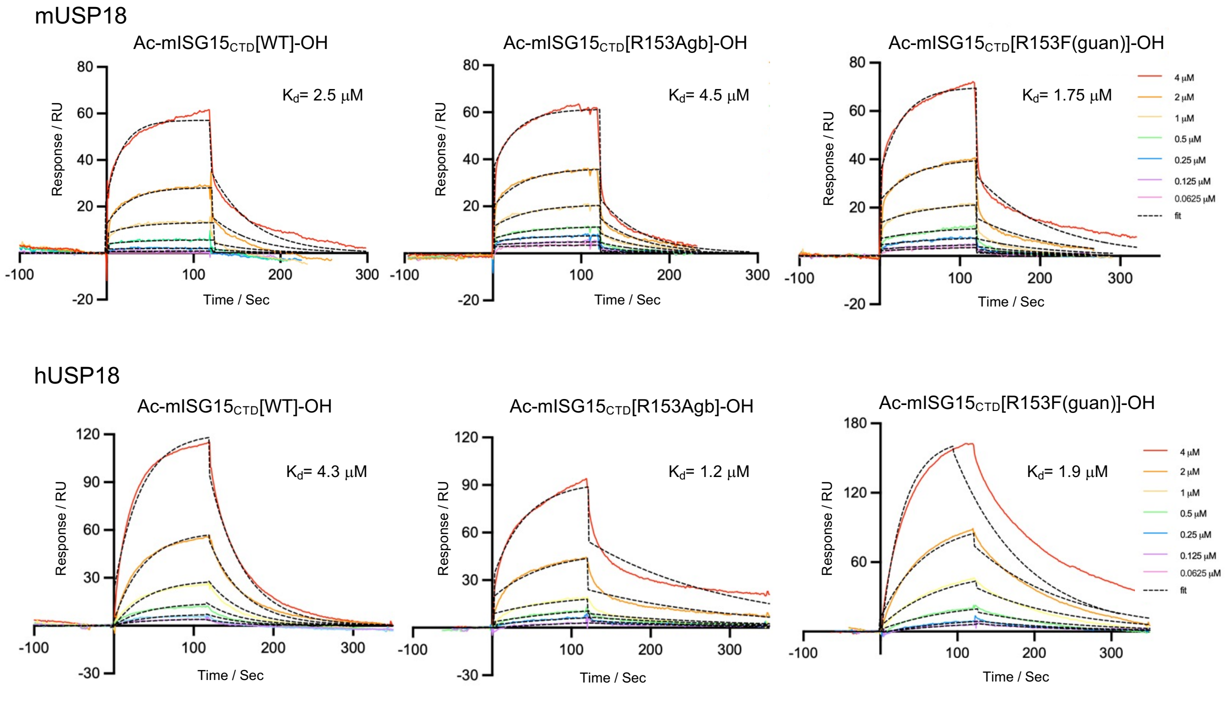


Figure S3. SPR analysis of the USP18-ISG15 interaction. Mouse or human USP18 was immobilized on Ni-NTA chips and association and dissociation of $mISG15_{CTD}$ variants were monitored.

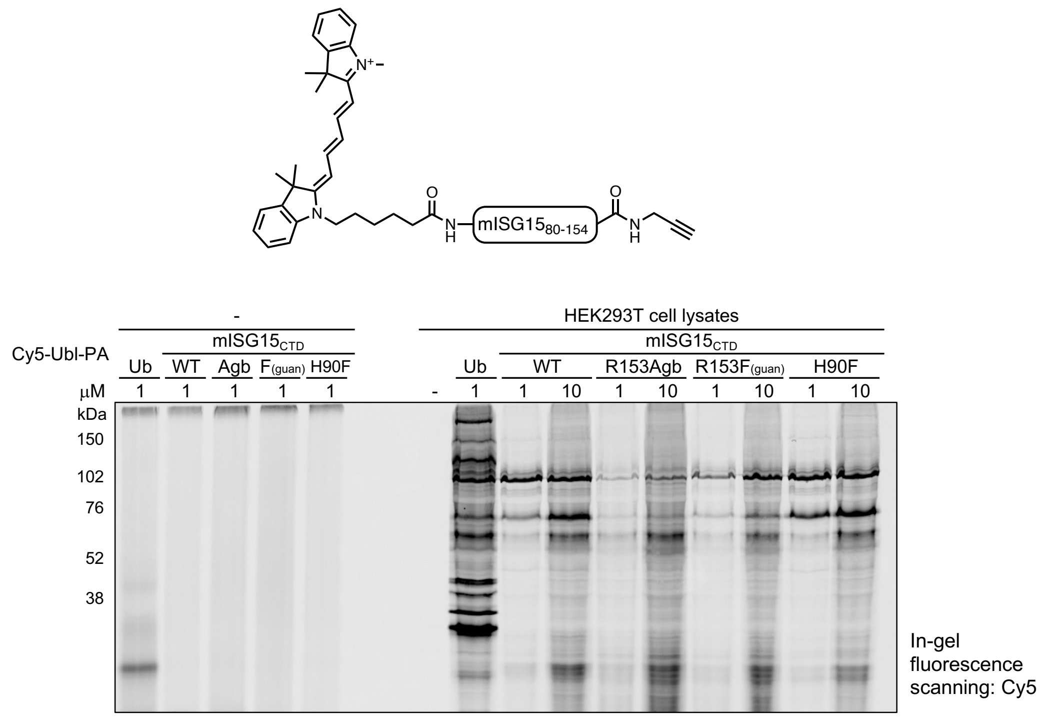


Figure S4. Selectivity of mISG15$_{CTD}$-based probes. HEK293T cell lysates were incubated with Cy5-mISG15$_{CTD}$-PA probes for 3 h at RT. Protein samples were analyzed by SDS-PAGE and in-gel fluorescence scanning for Cy5 signal.

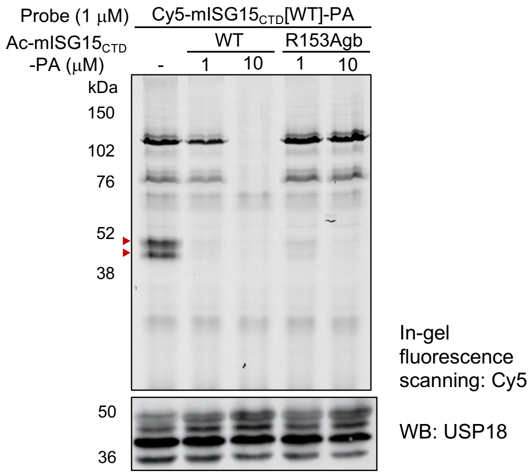


Figure S5. Selectivity of R153Agb mutant. USP18$^{WT}$-FLAG overexpressing HEK293T cell lysates were preincubated with Ac-mISG15$_{CTD}$-PA for 1 h followed by labeling with 1 μM of Cy5-mISG15$_{CTD}$[WT]-PA for 2 h at RT. Protein samples were analyzed by SDS-PAGE and in-gel fluorescence scanning for Cy5 signal. Red marks correspond to the expected molecular weight of USP18–probe conjugate.

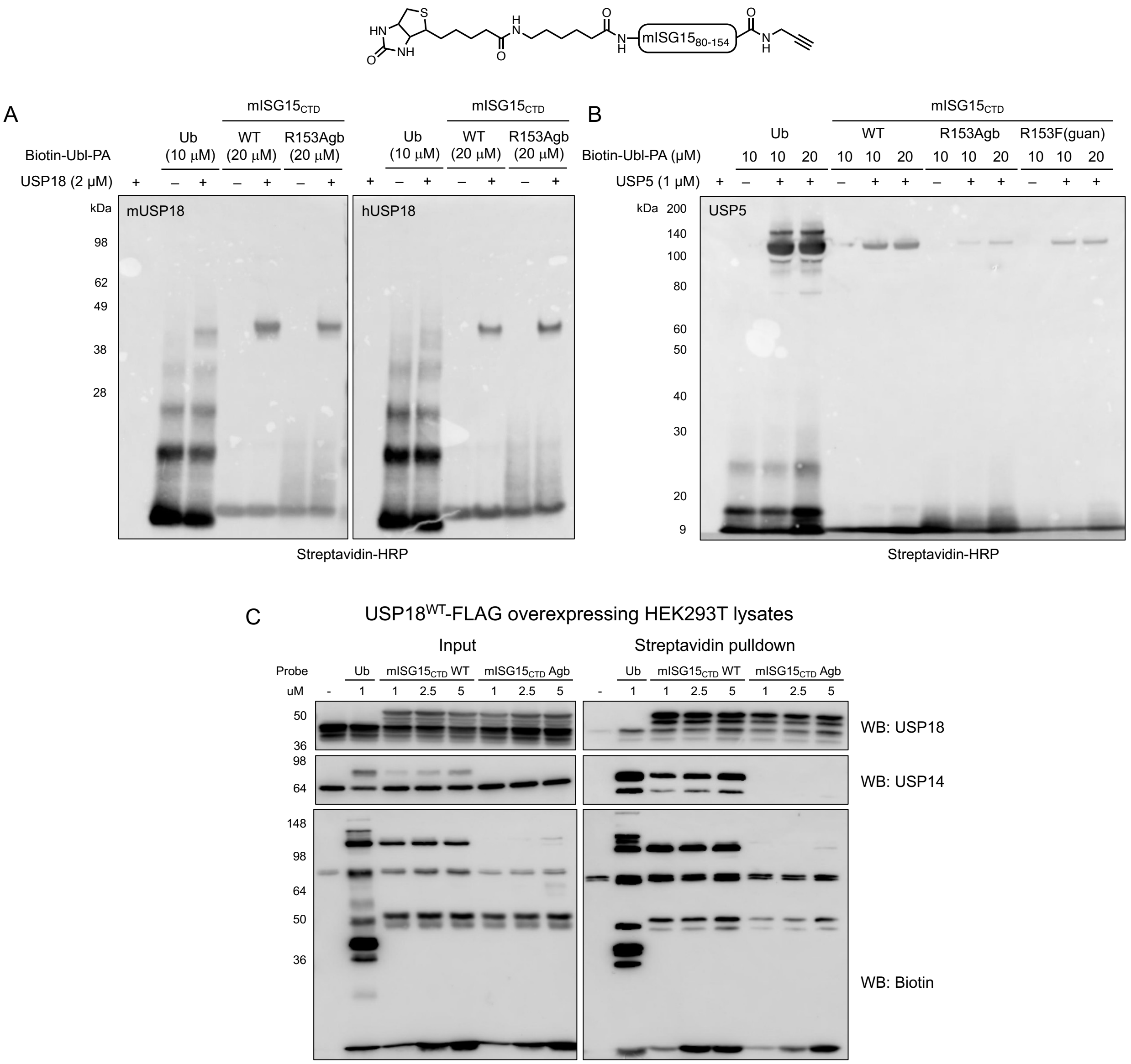


Figure S6. Reactivity and selectivity of Biotin-mISG15$_{CTD}$-based probes. (A-B) Recombinant human and mouse USP18 (2 μM) or USP5 (1 μM) was incubated with Biotin-mISG15$_{CTD}$-PA probes at indicated concentrations for 3 h at RT. Protein samples were analyzed by SDS-PAGE and far-western blotting using streptavidin-HRP. (C) USP18-FLAG overexpressing HEK293T cell lysates were incubated with Biotin-mISG15$_{CTD}$-PA probes at indicated concentrations for 3 h at RT. Biotinylated proteins were enriched by streptavidin pulldown and analyzed by SDS-PAGE and immunoblotting.

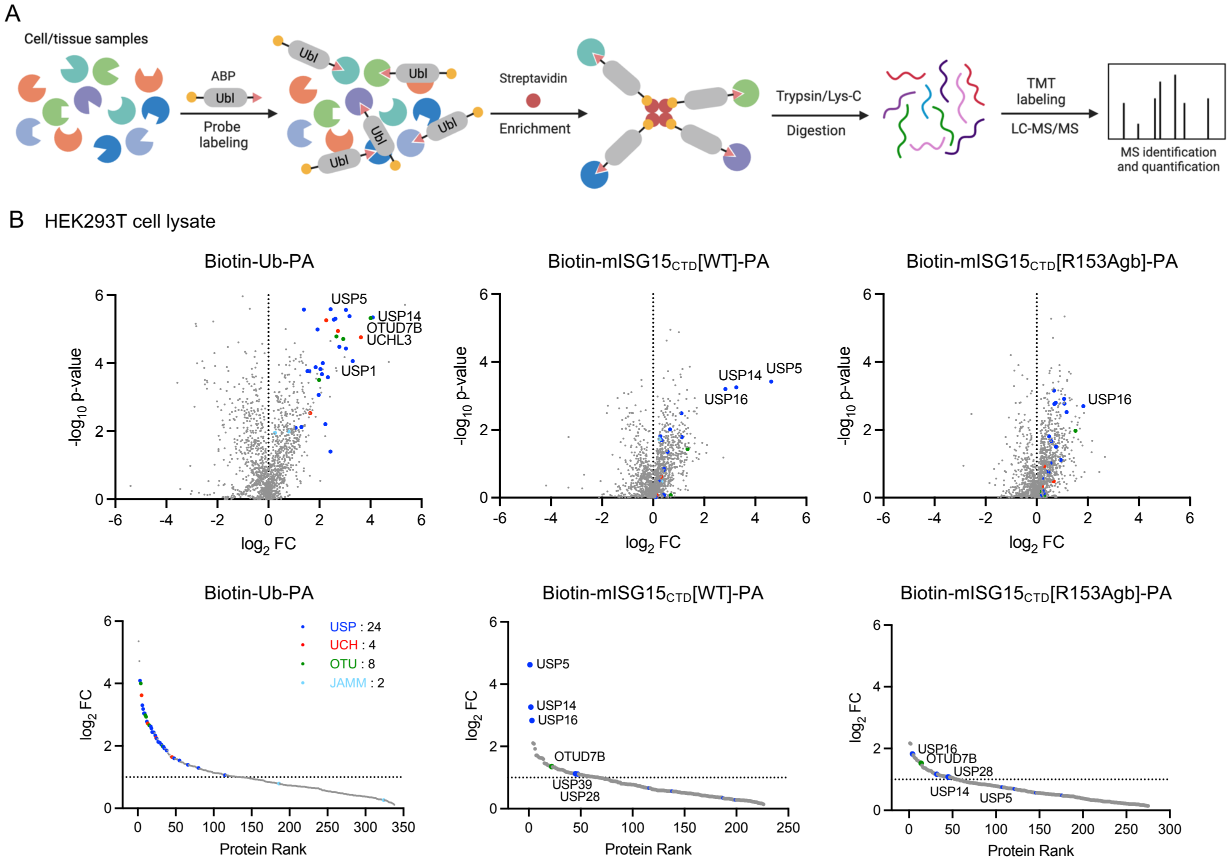


Figure S7. Activity-based protein profiling by Biotin-mISG15$_{CTD}$-PA probes. (A) Schematic workflow for identification of Ubl proteases in cell lysates. (B) Volcano (top) and protein rank (bottom) plots of quantitative proteomic analysis of streptavidin beads pulldowns after labeling of HEK 293T cell lysates by Biotin-Ub-PA, Biotin-mISG15$_{CTD}$[WT]-PA or Biotin-mISG15$_{CTD}$[R153Agb]-PA probes (1 µM, 2 h, RT) showing significantly enriched proteins ($log_2$ ratio > 1, p-value ≤ 0.05). DUBs are colored based on subfamilies.

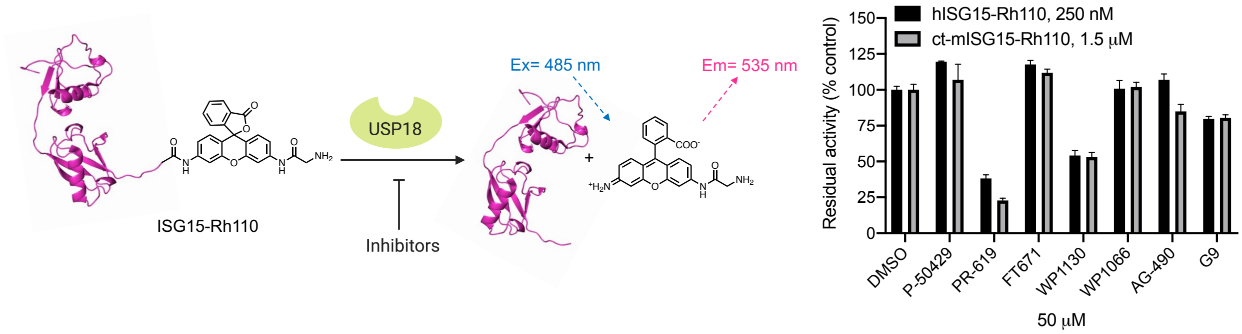


Figure S8. USP18 biochemical assay. (A) Schematic of ISG15-Rho screening assay. (B) Screening of known inhibitors of DUBs against mUSP18 (2.5 nM).

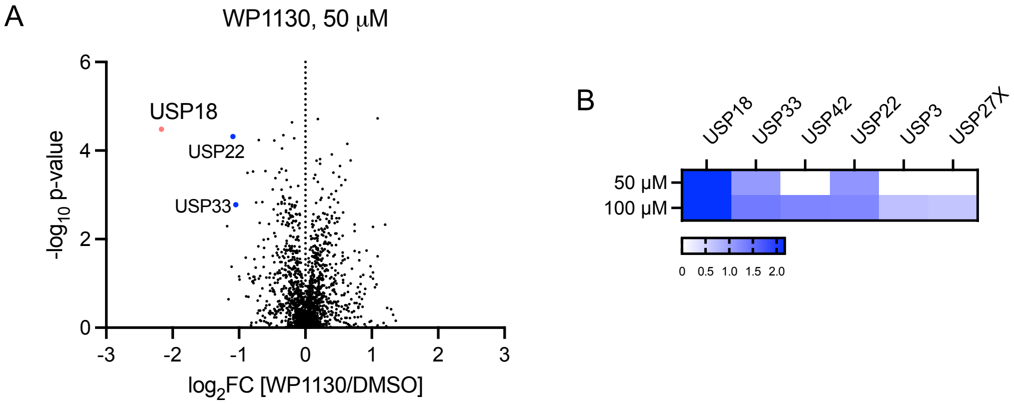


Figure S9. Competitive ABPP for DUB inhibitor screening. USP18$^{WT}$-FLAG overexpressing HEK293T cell lysates were preincubated with 50 μM of WP1130 for 2 h followed by labeling with a cocktail of probes (1 μM of Biotin-Ub-PA + 5 μM of Biotin-mISG15$_{CTD}$[R153Agb]-PA) for 3 h at RT. Protein samples were enriched by streptavidin, on-bead digested, and analyzed by LC-MS/MS after TMT labeling. (A) Volcano plots of quantitative proteomic analysis comparing samples treated with 50 μM of WP1130 to DMSO. USP18 is marked and other DUBs are colored based on subfamilies. (B) Heat map analysis of $\log_2$ fold change of MS intensity for enriched proteins comparing samples pretreated with 50 μM or 100 μM of WP1130 to DMSO samples.

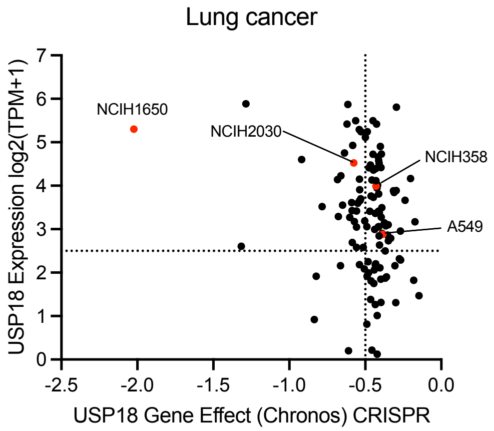


Figure S10. Analysis of lung cancer cell lines for USP18 expression and gene effect (CRISPR) from the Broad DepMap database. Red dots are cell lines used in this study.

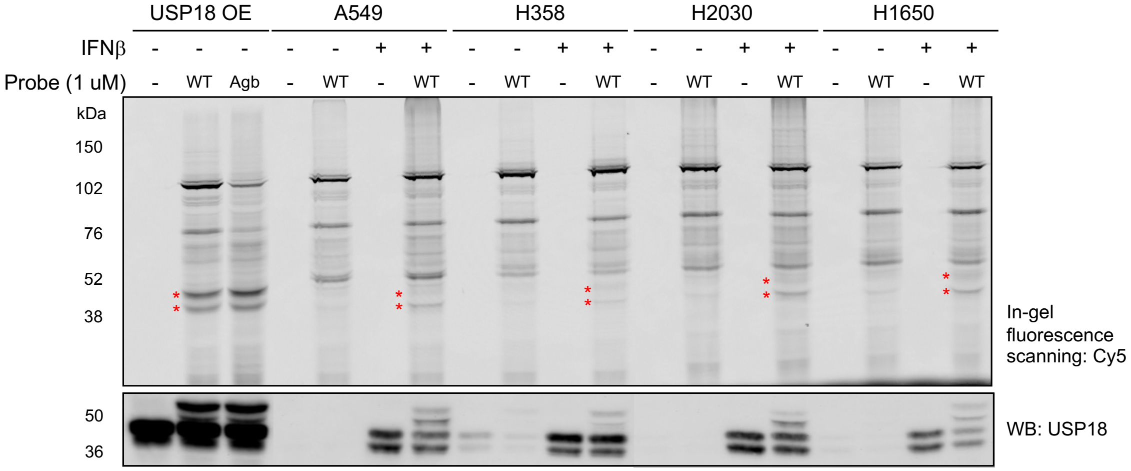


Figure S11. Gel-based ABPP in lung cancer cell lines. Each cell lysates were incubated with 1 μM of Cy5-mISG15$_{CTD}$[WT]-PA probes for 3 h at RT. Protein samples were analyzed by SDS-PAGE and in-gel fluorescence scanning for Cy5 signal. Red marks indicate the appearance of USP18–probe conjugates. Expression of USP18 was confirmed by western blotting.

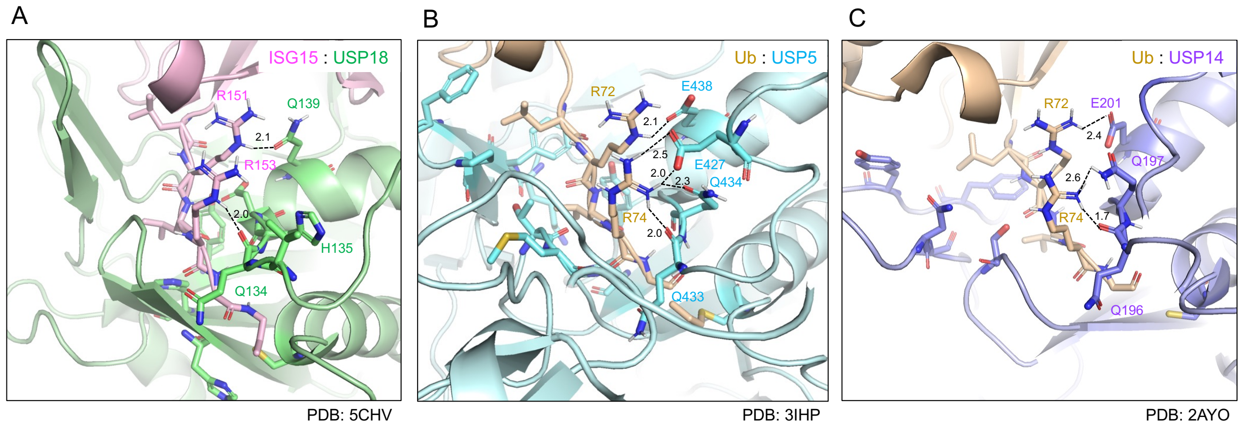


134QHDAAQ USP18
433QQDAQE USP5
196QQDANE USP14

Figure S12. Structure analysis of ISG15 cross-reactive DUBs for the interaction with Ubl C-termini. Close-up view of the area where the C-terminal tail of Ubl (LRGG motif) is guided to the catalytic center of DUBs with (A) mUSP18 (green) in complex with mISG15-PA (pink), (B) USP5 (cyan) in complex with Ub-PA (wheat), (C) USP14 (purple) in complex with Ub-Ald (wheat). Various residues important for interaction are shown as sticks.

### Materials and Methods

#### USP18 cloning, protein expression, and purification

Template plasmids for $His_6$-Mm.USP18 (46-368) and $His_6$-Hs.USP18 (16-372) with codon optimization for *Spodoptera frugiperda* (ATUM) were acquired and a TEV protease cleavage site (ENLYFQG) was added by PCR. Using Gateway BP Recombination (ThermoFisher Scientific), entry clones were generated, and both inserts were confirmed by Sanger sequencing. These entry clones were subcloned using Gateway LR Clonase (ThermoFisher Scientific) into pDest-795 (Addgene, 161881), a baculovirus expression vector that contains an N-terminal MBP tag. The expression clones were verified by restriction digestion and these plasmids were transformed into DE95 bacmid strain. Bacmid DNA was isolated by alkaline lysis and transposition of the desired MBP-tev-$His_6$ USP18 ORFS was confirmed by junction PCR.

The two USP18 clones were expressed following the insect cell culture as previously described.[1] The resulting cell pellets were stored at -80°C. Frozen cell pellets were thawed and resuspended in 100 mL of buffer per 1 L of culture in 20 mM HEPES, pH 7.3, 300 mM NaCl, 1 mM TCEP, and 1:500 (v/v) protease inhibitor cocktail. Lysis was performed using the M-110EH Microfluidizer (Microfluidics Corp., Westwood, MA) for 2 passes at 7,000 psi on ice. Lysates were then clarified by ultracentrifugation at 100,000×g for 30 min at 4°C and filtered through a Pall 250 ml Autofil 0.45 μm High Flow PES Bottle Top Filter (Thomas Scientific, Swedesboro NJ) and used immediately or stored at -80°C.

Clarified lysates were thawed, adjusted to 20 mM imidazole, and loaded onto IMAC columns equilibrated in IMAC equilibration buffer (EB) of 20 mM HEPES, pH 7.3, 300 mM NaCl, 1 mM TCEP with 20 mM imidazole. The columns were washed to baseline with EB with 20 mM imidazole and proteins eluted with a 20 column-volume (CV) gradient from 20 mM to 500 mM imidazole in EB. Elution fractions were analyzed by SDS-PAGE and Coomassie-staining. Positive fractions were pooled, and strep-TEV protease was added at 5% (v/v). The digest proceeded while dialyzing in a minimum buffer volume of 1:20 into EB for 2 h at room temperature followed by new buffer overnight at 4°C. The digested sample was purified via a second round of IMAC similar to the first round. The sample was loaded to the column equilibrated in EB and washed for 3 CV. Column flow through, wash, and elution bumps were collected as fractions. Columns were eluted with a 3 CV bump to EB with 35 mM imidazole, 3 CV bump wit EB with 70 mM imidazole, and a 2 CV bump in EB with 500 mM imidazole. After analysis of fractions by SDS-PAGE and Coomassie-staining, fractions containing the target of interest were pooled. The target protein was eluted in the 70 mM imidazole bump and dialyzed using 10k molecular weight cut-off snake skin into EB. Final samples were assayed for protein concentration, aliquoted in the final buffer (20 mM HEPES, pH 7.3, 300 mM NaCl, and 1 mM TCEP), and frozen.

#### Cell culture

HEK293T, HeLa cells were purchased from American Type Culture Collection (ATCC) and were cultured in Dulbecco's Modified Eagle Medium (DMEM, BTL, 112-013-101) containing 10% (v/v) fetal bovine serum (FBS, VWR International inc., 97068-901) supplemented with 100 units/mL penicillin-streptomycin (Sigma-Aldrich, P4333-100ML) and 2 mM L-glutamine (Gibco, 25030081) at 37°C in a humidified 5% $CO_2$ incubator. A549 cells were purchased from ATCC (CCL-185) and were cultured in Kaighn's Modification of Ham's F-12 Medium (F-12K, ATCC, 30-2004) containing 10% (v/v) FBS supplemented with 100 units/mL penicillin-streptomycin. H358, H2030 cells were provided by Ji Luo (NCI) and were cultured in Roswell Park Memorial Institute 1640 media (RPMI-1640, BTL, 112-024-101) containing 10% (v/v) FBS supplemented with 100 units/mL penicillin-streptomycin and 2 mM L-glutamine. H1650 cells were purchased from ATCC (CRL-5883) and were cultured in RPMI-1640 containing 10% (v/v) FBS supplemented with 100 units/mL penicillin-streptomycin and 2 mM L-glutamine.

**Stimulation of cells**

Cells were treated with 50 ng/mL of recombinant human IFN-β (PeproTech, 300-02BC) for the indicated times (48 h).

**Transfections**

pcDNA3.1-USP18 plasmids were purchased from GenScript. The plasmid was transformed into DH5α cells (Invitrogen, 18258012) The following day, a single transformed colony was used to inoculate 50 ml of LB medium (LB broth Lennox, Sigma-Aldrich, L3022-250g) containing 100 μg/ml of ampicillin (Sigma-Aldrich, A9518) and was incubated at 37°C overnight with agitation (250 rpm). A Midi prep (Qiagen, 12943) kit was used to isolate the plasmid for further experiments. HEK293T cells were grown in completed DMEM and maintained at 37°C with 5% $CO_2$. Each 100 mm cell culture dish was contained with 10 ml of complete DMEM media and transfected with 10 μg of overexpression plasmid with 30 μL Lipofectamine LTX (Invitrogen, 15338100) in 1 ml of Opti-MEM (Gibco, 31985070). After 48 h, cells were collected in PBS with 2 times of wash.

**Recombinant protein labeling assay**

Purified proteins were diluted in 25 mM Tris (pH 7.5), 150 mM NaCl, and 10 mM DTT. Purified Ac-mISG15$_{CTD}$-PA probes were diluted in 50 mM Tris (pH 7.5), 50 mM NaCl, and 5 mM DTT. The protein and probes were combined at a final concentration of 2 μM of USP18 or 1 μM of USP5 and 10 μM of probe and incubated for 3 h at room temperature. Reaction was stopped by addition of sample buffer and labeling was visualized by SDS-PAGE using NuPAGE 4-12% Bis-Tris gel with MES or MOPS running buffer and SYPRO Ruby staining.

**Activity-based protein profiling (ABPP) in lysates**

**Gel-based ABPP**

Using HEK293T cells transfected with plasmids genetically encoding full-length USP18$^{WT}$ and the catalytic mutant USP18$^{C64/65A}$, lysate was prepared 48 h after transfection in 50 mM Tris-HCl, 5 mM $MgCl_2$, 250 mM sucrose, and 1 mM DTT. 40 μg of lysate was treated with Cy5-Ubl-PA probe for 3 h at room temperature. Reaction was stopped by addition of reducing sample buffer and boiling. The samples were analyzed by SDS-PAGE and in-gel fluorescence scanning for Cy5 signal.

A549, H358, H1650, H2030 cells were prepared with or without IFN-β stimulation. Cell lysates were prepared in 50 mM Tris-HCl, 5 mM $MgCl_2$, 250 mM sucrose, and 1 mM DTT. 40 μg of total protein was treated with Cy5-Ubl-PA, WT, and Agb probe for 3 h at room temperature. Reaction was stopped by addition of reducing sample buffer and boiling. The samples were analyzed by SDS-PAGE and in-gel fluorescence scanning for Cy5 signal.

**Competitive gel-based ABPP**

40 μg of lysate was treated with either Ac-mISG15$_{CTD}$[WT]-PA or Ac-mISG15$_{CTD}$[R153Agb]-PA at a concentration of 1 μM/10 μM and allowed to incubate for 1 h at room temperature. Samples were then chased with 1 μM Cy5- mISG15$_{CTD}$[WT]-PA for 2 h at room temperature, and the reaction was stopped by addition of reducing sample buffer and boiling. The samples were analyzed by SDS-PAGE and in-gel fluorescence scanning for Cy5 signal.

**Western blotting**

To validate USP18 expression and labeling following gel-based ABPP, SDS-PAGE samples were transferred onto a nitrocellulose membrane (Bio-Rad). The membranes were blocked with 2% (w/v) BSA for 1 h, washed with TBST (containing 0.1% (v/v) Tween 20), and then incubated with anti-USP18 (Cell Signaling, 4813) primary antibody overnight at 4°C. After additional washing with TBST, membranes were probed with a horseradish peroxidase-conjugated secondary antibody (Cell Signaling, 7074) for 1 h at room temperature, followed by ECL detection (SignalFire ECL Reagent, Cell Signaling). The protein bands were observed using ImageQuant 800 imaging system (Cytiva). For far-western blotting of probe-labeled recombinant enzymes, the nitrocellulose membrane was incubated with streptavidin-horseradish peroxidase antibody (Cell Signaling, 3999) for 1 h at room temperature and subsequently imaged following ECL detection.

**Quantitative MS-based ABPP**

HEK 293T cells were lysed (50 mM Tris pH 8.0, 150 mM NaCl, 5 mM $MgCl_2$, 0.5 mM EDTA, 0.5% NP-40, 10% glycerol, 1 mM TCEP, protease inhibitor (cOmplete™ Mini Protease Inhibitor Cocktail, Roche)), clarified by centrifugation, and then diluted to a concentration of 2 mg/mL. 500 μL aliquots were incubated with 50 or 100 μM of the WP or DMSO for 2 h at room temperature. Subsequently, 1 μM of Ub or EY-2-137(Agb) probe were treated for 3 h at room temperature. To Quench the reaction, 500 μL of 0.2% SDS, 0.5% NP-40 in PBS were added to cell lysate (Final 0.1% SDS, 0.25% NP-40). 100 μL of high-capacity streptavidin agarose resin slurry (ThermoFisher Scientific) was added to low-binding microcentrifuge tubes (Sorenson) and washed (3 × PBS) and 200 μL of 0.2% NP-40/PBS added before mixing with cell lysate. Protein samples were added to the streptavidin beads and incubated at room temperature for 1 h with end-to-end rotation. After incubation, beads were washed (4 × 0.2% NP-40/PBS, 4 × PBS, 4 × dd$H_2O$, 1 × HEPES) by centrifugation (500 × g, 1 min, 4 ° C) with Micro Bio-Spin™ Chromatography Columns (Bio-rad). After the final wash, the supernatant was removed, 50 μL of 100 mM of HEPES pH 7.4 was added, and the beads were flash frozen in liquid $N_2$ and stored at -80 °C.

**Identification of probe-labeled proteins by mass spectrometry**

The frozen streptavidin beads were thawed at room temperature and each were added with 200 μL of 1:1:1:1 100 mM HEPES/lysis buffer/reducing solution/alkylating solution/ from the EasyPep™ MS Sample Prep Kit (Thermo, A40006) containing 3 μg Trypsin/LysC protease mix (Thermo, A40007). The samples were incubated overnight at 37 °C with shaking and 200 μL aliquot from each sample was obtained. TMTpro 18-plex reagents (125 μg, Thermo, A52045) were dissolved in 20 μL acetonitrile (ACN) and added into each sample, followed by incubation at 25 °C for 1 hr. Samples were quenched with 50 μL of 5% hydroxylamine + 20% formic acid (FA) for 15 mins room temperature. The samples were pooled and cleaned up using EasyPep Mini columns (Thermo, A40006) by following the manufacturer's protocol. The eluted peptides were dried under vacuum in a speed-vac and stored in -20 °C.

The dried peptides were resuspended in 50 μL of 0.1% FA and loaded 5 µL twice onto a Dionex U3000 RSLC in front of a Orbitrap Eclipse (Thermo) equipped with an EasySpray ion source Solvent A consisted of 0.1% FA in water and Solvent B consisted of 0.1% FA in 80% ACN. Loading pump consisted of Solvent A and was operated at 7 μL/min for the first 6 minutes of the run then dropped to 2 μL/min when the valve was switched to bring the trap column (Acclaim™ PepMap™ 100 C18 HPLC Column, 3 µm, 75 µm I.D., 2 cm, PN 164535) in-line with the analytical column EasySpray C18 HPLC Column, 2 µm, 75 µm I.D., 25 cm, PN ES902). The gradient pump was operated at a flow rate of 300 nL/min and each run used a linear LC gradient of 5-7%B for 1 min, 7-30% B for 134 min, 30-50% B for 35 min, 50-95% B for 4 min, holding at 95% B for 7 min, then re-equilibration of analytical column at 5% B for 17 min. All MS injections employed the TopSpeed method with four FAIMS compensation voltages (CVs) and a 0.75 second cycle time for each CV (3 second cycle time total) that consisted of the following: Spray voltage was 2200V and ion transfer temperature of 300 °C. MS1 scans were acquired in the Orbitrap with resolution of 120,000, AGC of 4e5 ions, and max injection time of 50 ms, mass range of 375-1600 m/z; MS2 scans were acquired in the Orbitrap using TurboTMT method with resolution of 15,000, AGC of 1.25e5, max injection time of 22 ms, HCD energy of 38%, isolation width of 0.4 Da, intensity threshold of 2.5e4 and charges 2-6 for MS2 selection. Advanced Peak Determination, Monoisotopic Precursor selection (MIPS), and EASY-IC for internal calibration were enabled and dynamic exclusion was set to a count of 1 for 15sec. The only difference in the methods was the

CVs used: one method used CVs of -45, -55, -65, -75 and the second used CVs of -50, -60, -70, -80.

**Database search and data post-processing**

Both injections for each sample were pooled together as fractions and all MS files were searched with Proteome Discoverer 2.4 using the Sequest node. Data was searched against the Uniprot Human database from Feb 2020 using a full tryptic digest, 2 max missed cleavages, minimum peptide length of 6 amino acids and maximum peptide length of 40 amino acids, an MS1 mass tolerance of 10 ppm, MS2 mass tolerance of 0.02 Da, variable oxidation on methionine (+15.995 Da) and fixed modifications of carbamidomethyl on cysteine (+57.021), TMTpro (+304.207) on lysine and peptide N-terminus. Percolator was used for FDR analysis and TMTpro reporter ions were quantified using the Reporter Ion Quantifier node and normalized on total peptide intensity of each channel. TMTpro channel assignment for conditions can be found in supplementary tables.

**Combinatorial substrate library synthesis**

A combinatorial tetrapeptide fluorogenic substrate library was synthesized on a solid support according to published protocols.[2, 3] The library consisted of two tetrapeptide sub-libraries. Each of the sub-libraries was synthesized separately, and the general synthetic procedure is described for the P3 sub-library as an example. In the first step, Fmoc-ACC-OH (25 mmol, 2.5 eq.) was attached to the Rink amide resin (13.5 g) using coupling reagents: HOBt (25 mmol, 2.5 eq.) and DICI (25 mmol, 2.5 eq.) in DMF. After 24 h, the Fmoc protecting group was removed with 20% piperidine in DMF. In the next step, Fmoc-Gly-OH (25 mmol, 2.5 eq.) was coupled using HATU (25 mmol, 2.5 eq.) and 2,4,6-collidine (25 mmol, 2.5 eq.) in DMF. Then, Fmoc-Gly-OH (25 mmol, 2.5 eq.) was attached to the $H_2N$-Gly-ACC-resin using HOBt and DICI (25 mmol, 2.5 eq.) as coupling reagents. After glycine coupling, the Fmoc group was removed (20% piperidine in DMF), and the resin was washed with DCM and MeOH and dried over $P_2O_5$. Then, the dried resin was divided into 138 portions. To each portion of the $H_2N$-Gly-Gly-ACC-resin, natural or unnatural amino acids were attached, and the Fmoc protecting group was removed (20% piperidine in DMF). To provide an equimolar substitution of each natural amino acid in the P4 position, an isokinetic mixture of Fmoc-protected amino acids was utilized. The last two steps of P3 sub-library synthesis included N-terminal acetylation (solution of AcOH, HBTU and DIPEA) and cleavage of peptides from the resin using a TFA:$H_2O$:TIPS (95:2.5:2.5, % v/v/v) mixture. Finally, the sub-library was precipitated in $Et_2O$, dissolved in a mixture of acetonitrile and water, lyophilized, and dissolved in biochemical grade DMSO at a concentration of 20 mM. The obtained sub-library was used for kinetic studies without further purification. The P4 sub-library was synthesized in the same manner.

**Substrate library screening**

All screenings were performed using a spectrofluorometer (Molecular Devices Spectramax Gemini XPS) in 96-well plates (Corning). The release of ACC was monitored continuously for 40 min at the appropriate wavelength (Ex = 355 nm, Em = 460 nm). For the assay, 0.5 μL of substrate in DMSO was used with 49.5 μL of enzyme. The enzyme was incubated in assay buffer (50 mM Tris, 100 mM NaCl, 1 mM TCEP, 0.01% BSA, 0.01% Triton X-100, pH 7.5) for 30 min at 37°C before addition to the substrates on a plate. The final substrate concentration in each well during the assays was 200 μM for combinatorial P3 and P4 sub-libraries. The enzyme concentration was 10 μM for mUSP18. The linear part of each progress curve was used to determine the substrate hydrolysis rate. Substrate specificity profiles were established by setting the highest value of relative fluorescence unit per second (RFU/s) from each library position as 100% and adjusting other values accordingly.

**Surface plasmon resonance (SPR) analysis**

Surface plasmon resonance binding analysis was performed on a BIAcore 3000 (GE Healthcare) instrument. Mouse and human USP18 were immobilized via the N-terminal $His_6$ tag on NTA chip (Cytiva). Protein binding analysis was performed at 25°C in 50 mM HEPES, 150 mM NaCl, 0.05 mM EDTA, 0.5 mM DTT and 0.01% P20, pH 7.5, with a flow rate of 20 μL/min. Stock solutions of ubiquitin, ISG15 and mouse ISG15 variants were diluted in running buffer. Binding traces were analyzed with BIAevaluation 4.0 software (GE Healthcare) and fitted with a Langmuir 1:1 binding model including a drift of the baseline.

**Preparation of ISG15-based probes**

*Ac-mISG15$_{CTD}$-PA*

The C-terminal domain of mouse ISG15 was synthesized using solid-phase peptide synthesis (SPPS) to yield the sequence Ac-LSILVRNERGHSNIYE VFLTQTVDTLKKKVS QREQVHEDQF WLSFE GRPMEDKELLGEYGLKPQ**S**TVIKHLRLRG (wherein the underlined portions denote positions where pseudoproline dipeptides were used, and the bolded residue in position 144 indicates a Cys→Ser change from the native mISG15 sequence).[4] For the R153Agb and R153F(guan) mutant sequences, non-canonical amino acids were manually coupled to pre-loaded Fmoc-Gly Tentagel R Trt resin (RAPP Polymere 0.17 mmol/g) prior to automated synthesis. All sequences were synthesized at a 25 μmol scale in *N*-methylpyrrolidone (NMP) with PyBOP (4 eq., 50 μmol, 26 mg) and diisopropylethylamine (8 eq., 100 μmol, 17.5 μL) using Tetras synthesizer (Thuramed) with Fmoc-based SPPS, and reaction progress was periodically monitored by microcleavage. Upon completion, 12.5 μmol of the protected peptide was cleaved from the resin using 1,1,1,3,3,3-hexafluoroisopropanol in DCM (20% (v/v), 2 × 30 min.) and the cleavage mixture was removed by evaporation, after which the cleaved peptide was coevaporated with 1,2-dichloroethane. After drying under vacuum, the resulting peptide was resuspended in 3 mL dry DCM and stirred overnight with PyBOP (4 eq., 50 μmol, 26 mg), triethylamine (4 eq., 50 μmol, 7 μL), and propargylamine (10 eq., 125 μmol, 8 μL). All solvents were evaporated, and the peptide mixture was then fully deprotected by stirring in TFA/$H_2O$/TIPS/phenol (90:5:2.5:2.5 (v/v/v/v)) for 3.5 h. The crude, deprotected peptide was precipitated from the cleavage mixture through addition of chilled ether:*n*-heptane = 3:1 solution (v/v) and spun down. The pellet was allowed to air dry, and then resuspended in $H_2O$:ACN:AcOH = 65:25:10 (v/v/v) and lyophilized. The crude product was then purified by RP-HPLC on a Teledyne ACCQ Prep HP 150 with a C18 column (Redisep). The mobile phases were; A, 0.1% TFA in $H_2O$; B, 0.1% TFA in ACN. At a flow rate of 20 mL/min, a gradient of 20-80% B over 30 min was used. Fractions containing the pure product were combined and lyophilized.

*Biotin-mISG15$_{CTD}$-PA*

The protocol for SPPS was followed exactly as specified above. 12.5 μmol of resin was swelled in NMP for 30 min prior to Fmoc-deprotection in piperidine/NMP (20% (v/v), 30 min). The resin was then allowed to shake overnight with biotin-X-NHS (4 eq., 50 μmol, 23 mg) and diisopropylethylamine (16 eq., 200 μmol, 35 μL) in 3 mL NMP. Biotinylated polypeptide was then cleaved from the resin using 1,1,1,3,3,3-hexafluoroisopropanol in DCM (20% (v/v), 2 × 30 min), and all steps for C-terminal propargylation were followed as described above.

*Cy5-mISG15$_{CTD}$-PA*

The protocol for SPPS was followed exactly as specified above. 12.5 μmol of resin was swelled in NMP for 30 min prior to Fmoc-deprotection in piperidine/NMP (20% (v/v), 30 min). The resin was then washed with NMP and DMF, successively. The resin was then allowed to shake overnight in the dark with Cy5-NHS ester (1 eq., 12.5 μmol, 8 mg) and diisopropylethylamine (16 eq., 200 μmol, 35 μL) in 3 mL DMF. Cy5-coupled polypeptide was then cleaved from the resin using 1,1,1,3,3,3-hexafluoroisopropanol in DCM (20% (v/v), 2 × 30 min), and all steps for C-terminal propargylation were followed as described above.

*Ac-mISG15$_{CTD}$-OH*

The protocol for SPPS was followed exactly as specified above. The completed peptide was then cleaved from the resin and fully deprotected simultaneously, through treatment with TFA:$H_2O$:TIPS:phenol = 90:5:2.5:2.5 (v/v/v/v) for 3.5 h. All precipitation and purification steps were followed as specified for the C-terminal propargylated probes above.

## LC-MS

*Ac-mISG15$_{CTD}$[WT]-OH*

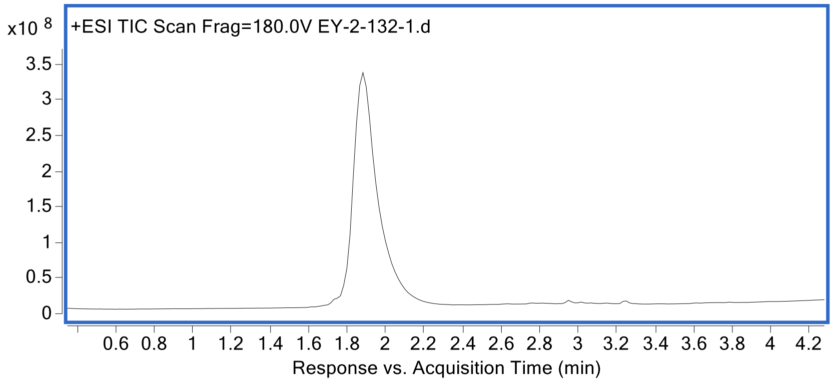


HRMS (ESI): m/z [M] calc. for $C_{399}H_{638}N_{114}O_{118}S$: 8952.2330, found 8952.71.

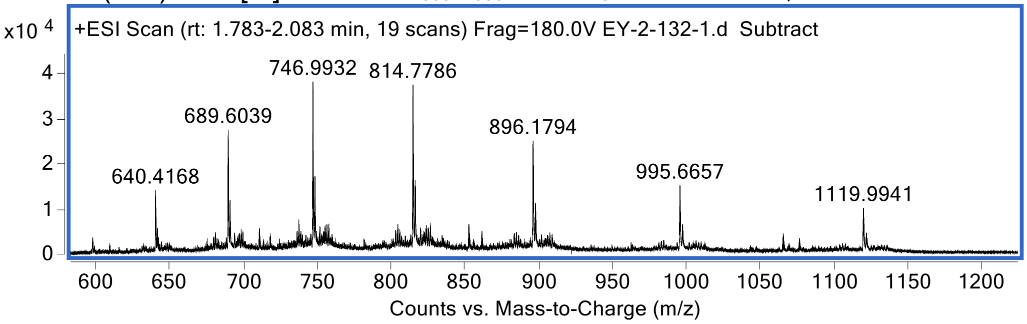


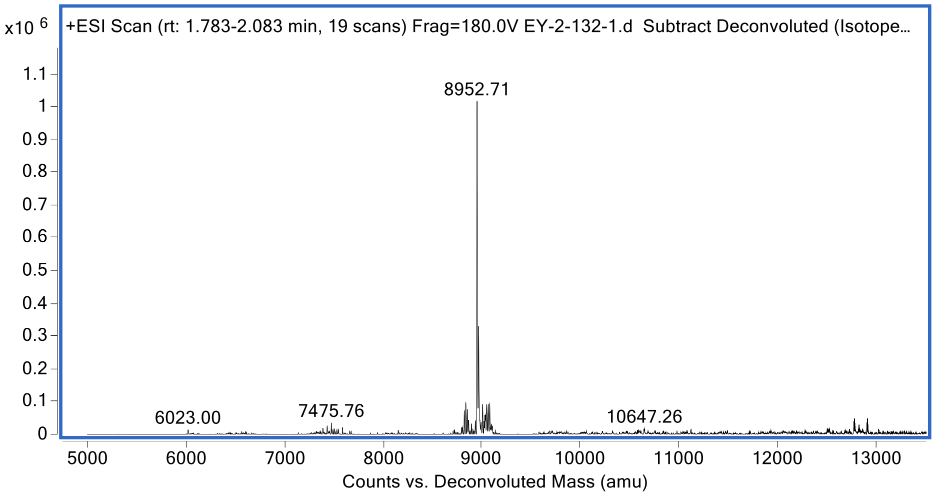

*Ac-mISG15$_{CTD}$[R153Agb]-OH*

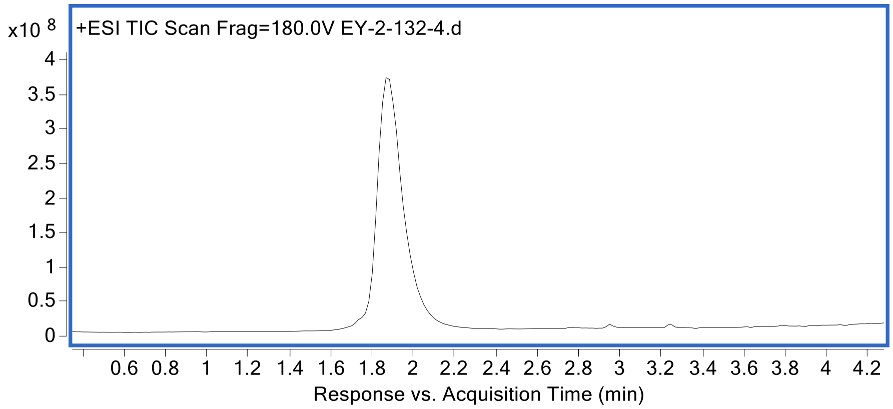


HRMS (ESI): m/z [M] calc. for $C_{398}H_{636}N_{114}O_{118}S$: 8938.2060, found 8938.54.

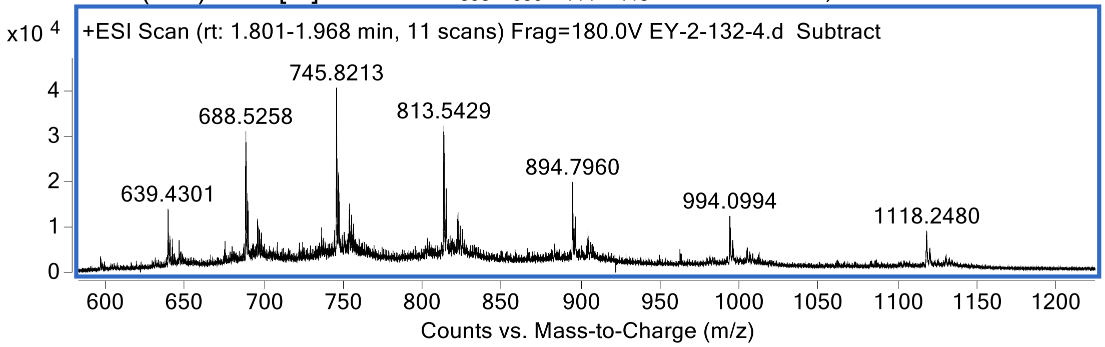


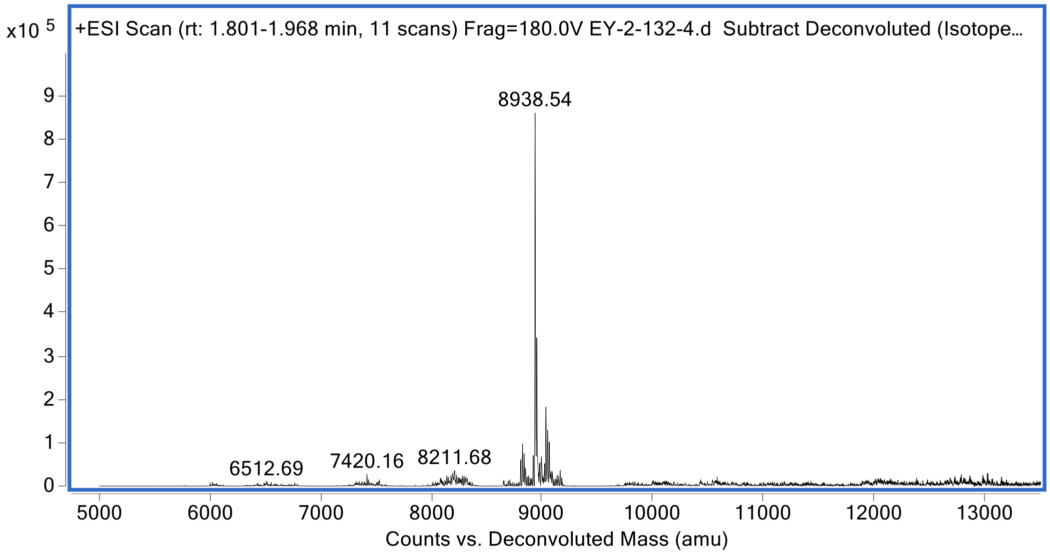

*Ac-mISG15$_{CTD}$[R153F(guan)]-OH*

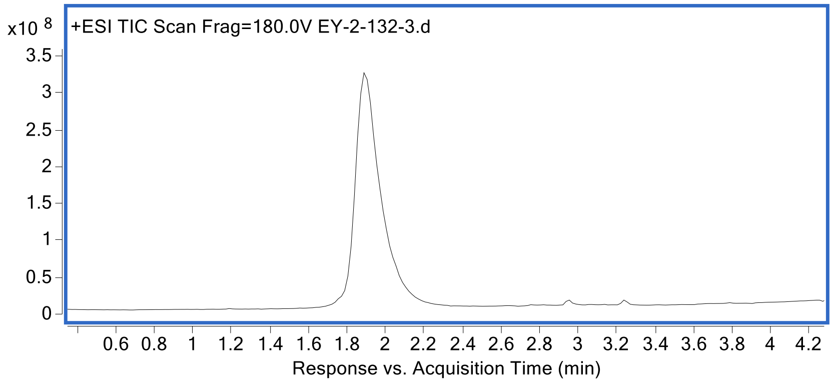


HRMS (ESI): m/z [M] calc. for $C_{403}H_{638}N_{114}O_{118}S$: 9000.2770, found 9000.68.

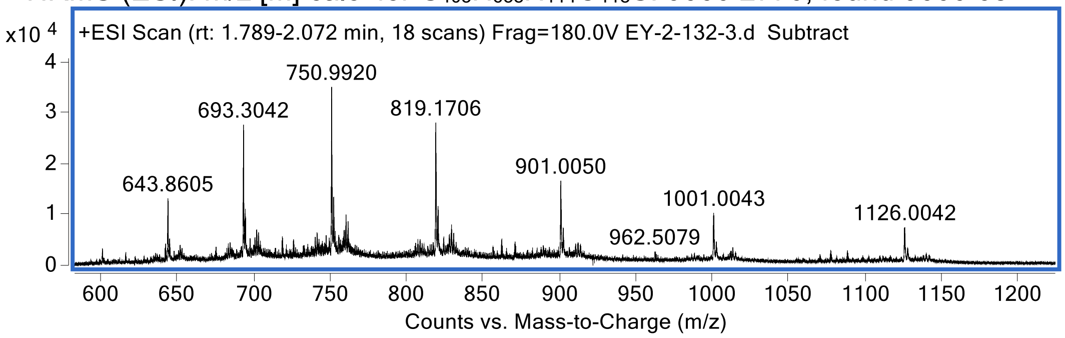


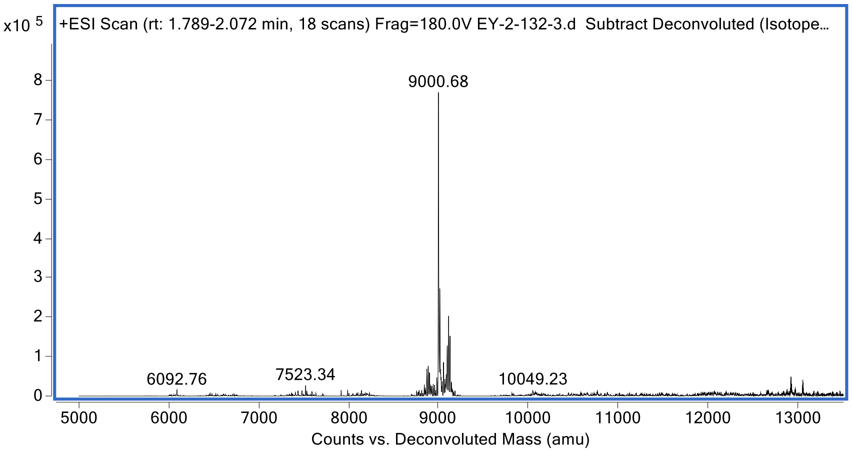

*Ac-mISG15$_{CTD}$[WT]-PA*

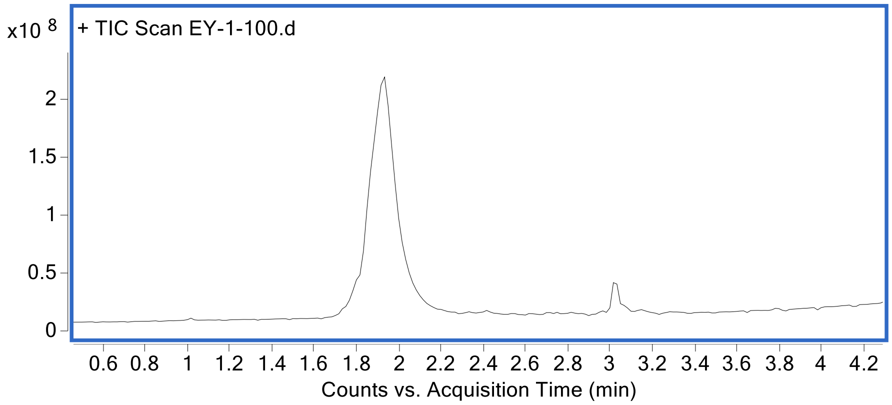


HRMS (ESI): m/z [M] calc. for $C_{400}H_{638}N_{114}O_{116}S$: 8932.25, found 8932.66.

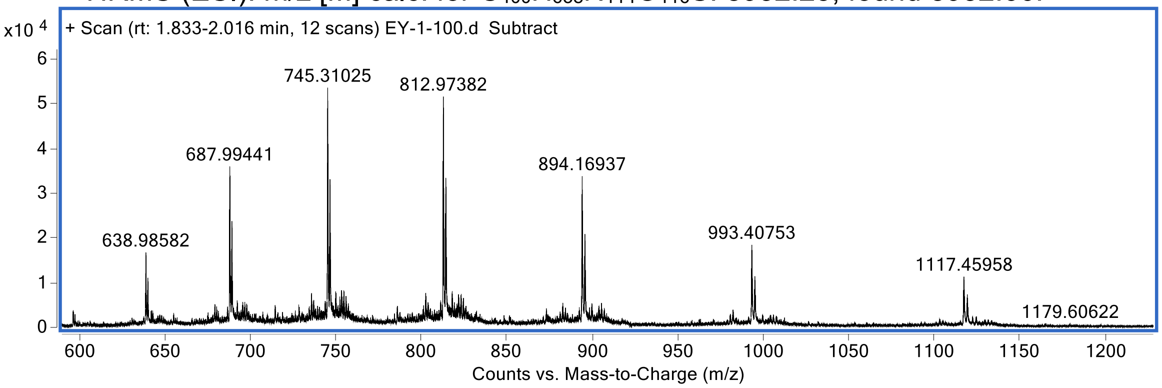


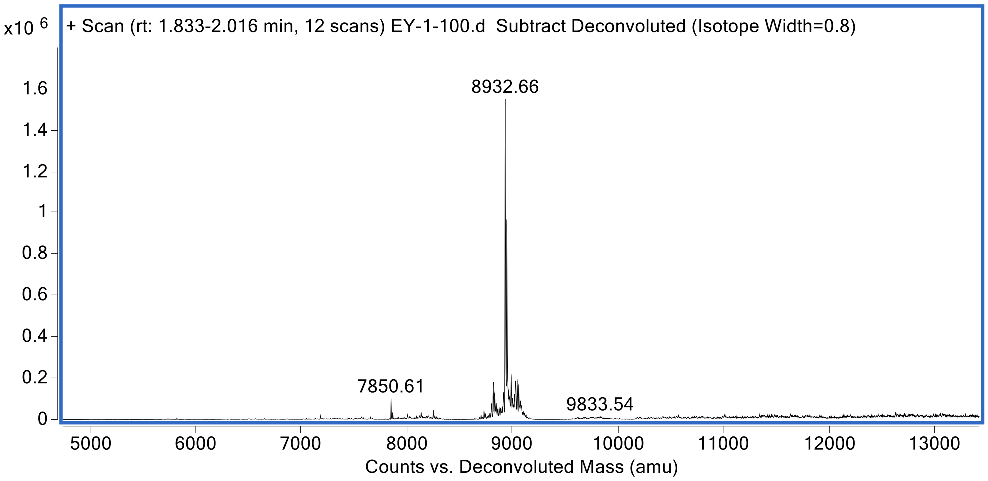

*Ac-mISG15$_{CTD}$[R153Agb]-PA*

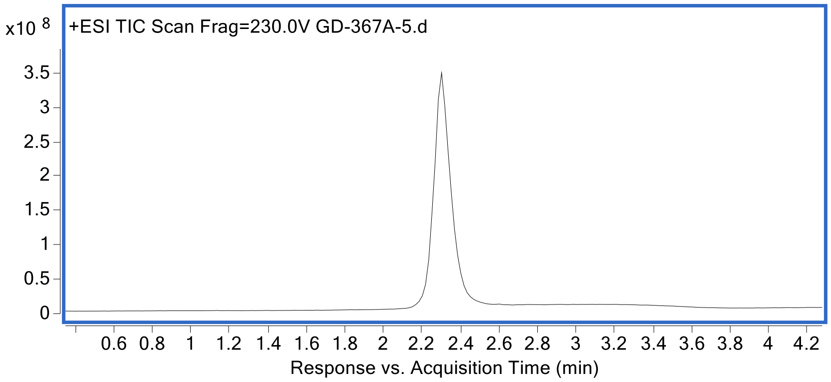


HRMS (ESI): m/z [M] calc. for $C_{399}H_{636}N_{114}O_{116}S$: 8918.2190, found 8934.64 (+16.421)

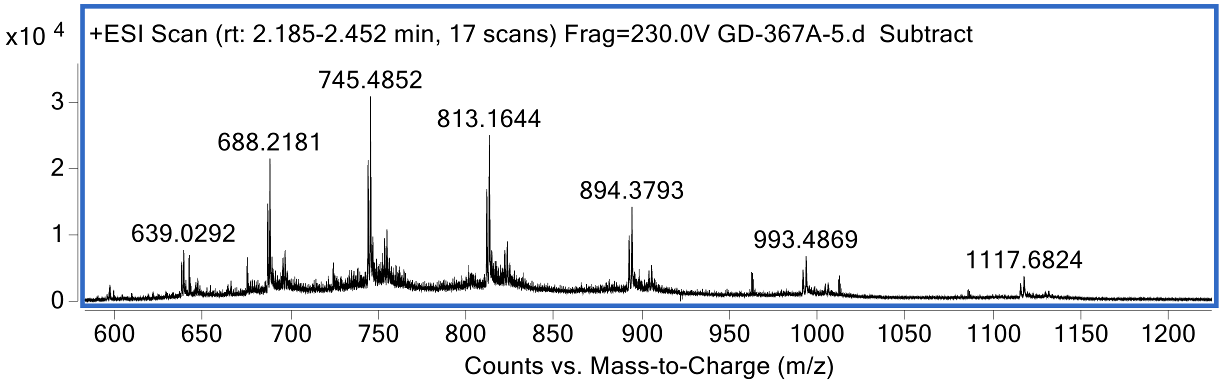


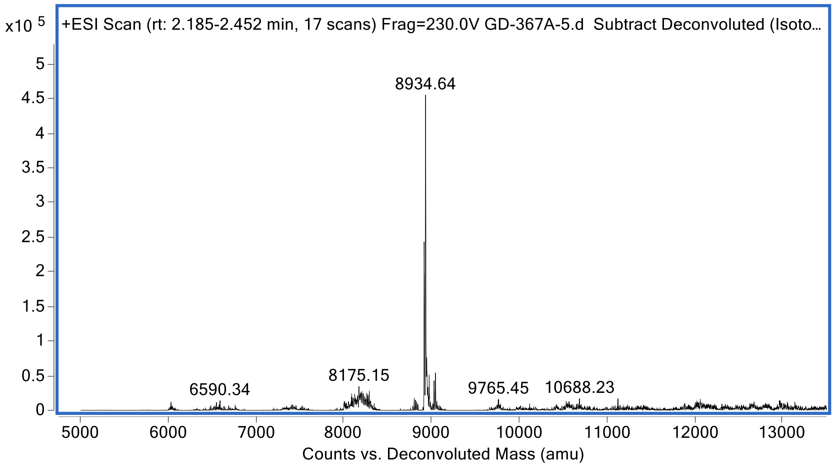

*Ac-mISG15$_{CTD}$[R153hR]-PA*

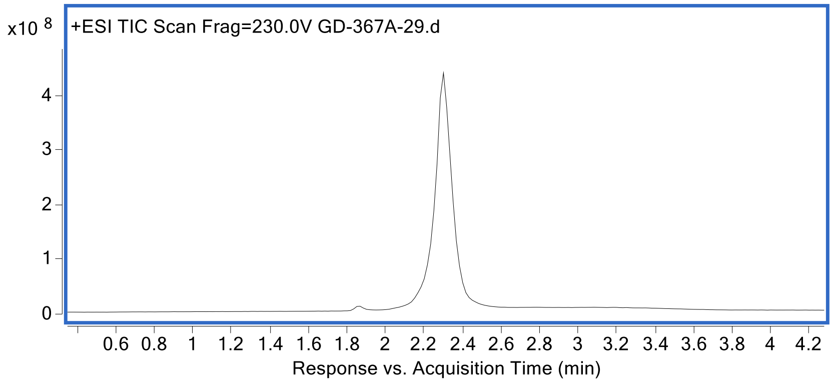


HRMS (ESI): m/z [M] calc. for $C_{401}H_{640}N_{114}O_{116}S$: 8946.273, found 8962.71 (+16.437)

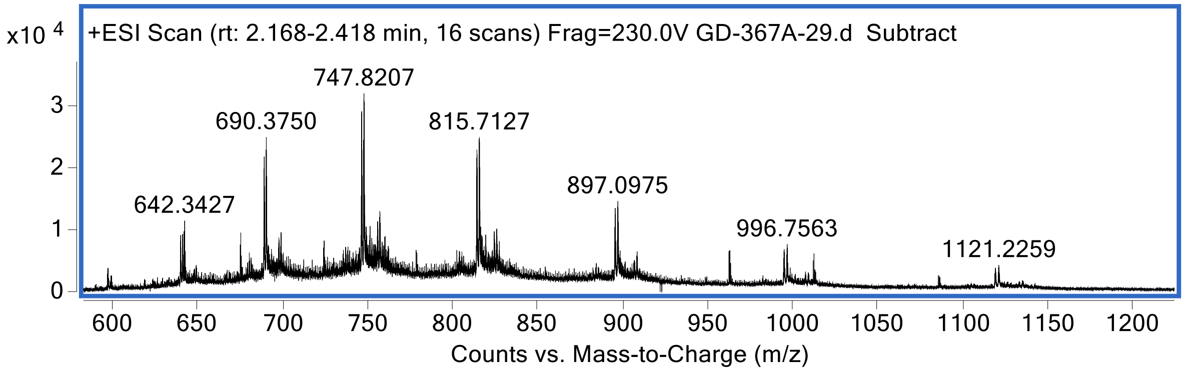


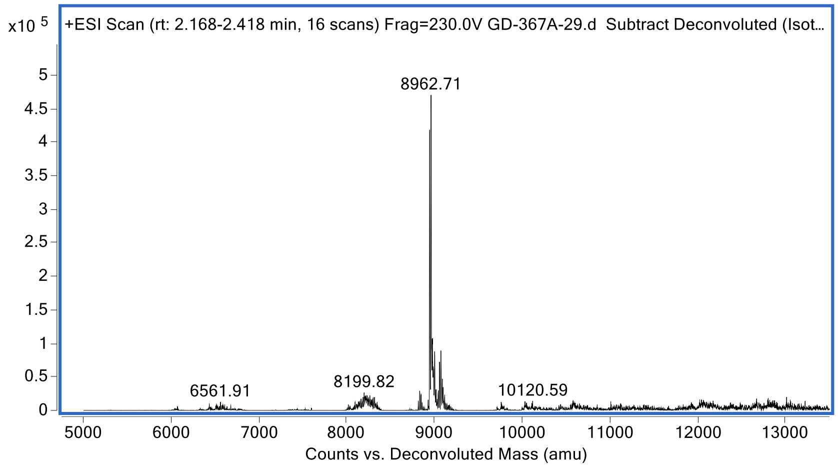

*Ac-mISG15$_{CTD}$[R153F(guan)]-PA*

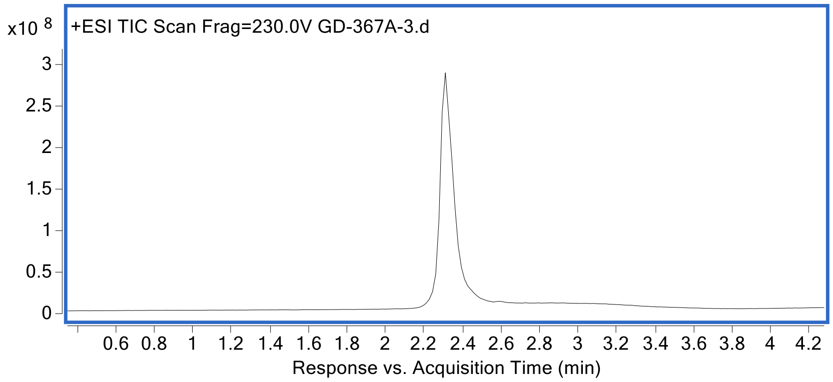


HRMS (ESI): m/z [M] calc. for $C_{404}H_{638}N_{114}O_{116}S$: 8980.2900, found 8980.60.

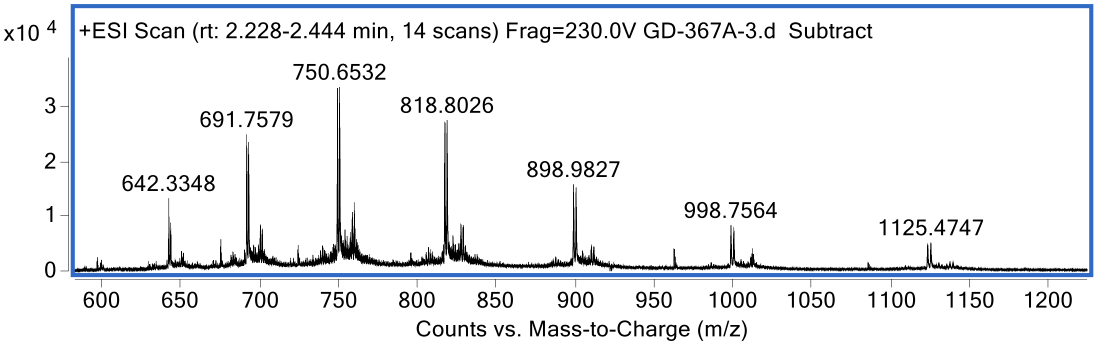


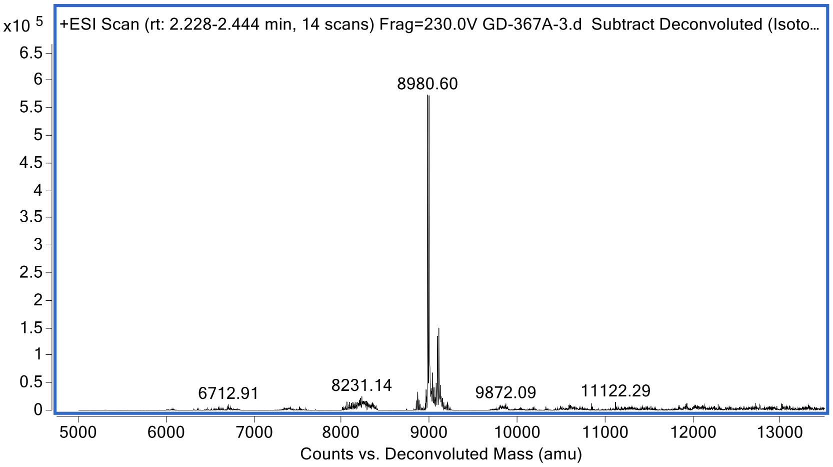

*Cy5-mISG15$_{CTD}$[WT]-PA*

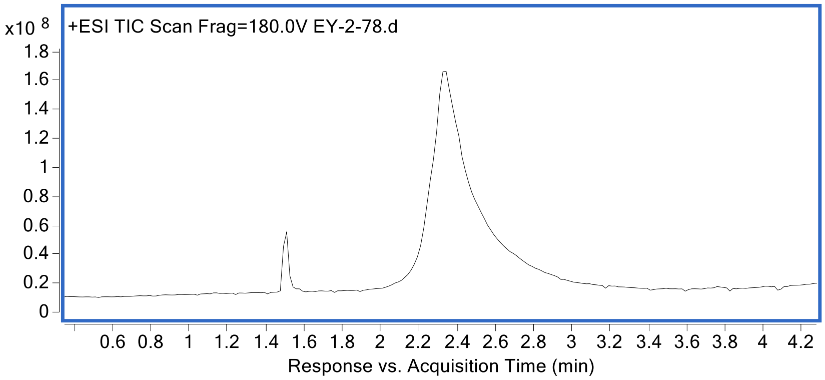


HRMS (ESI): m/z [M] calc. for $C_{430}H_{673}N_{116}O_{116}S$: 9355.8695, found 9370.84. (+14.9705)

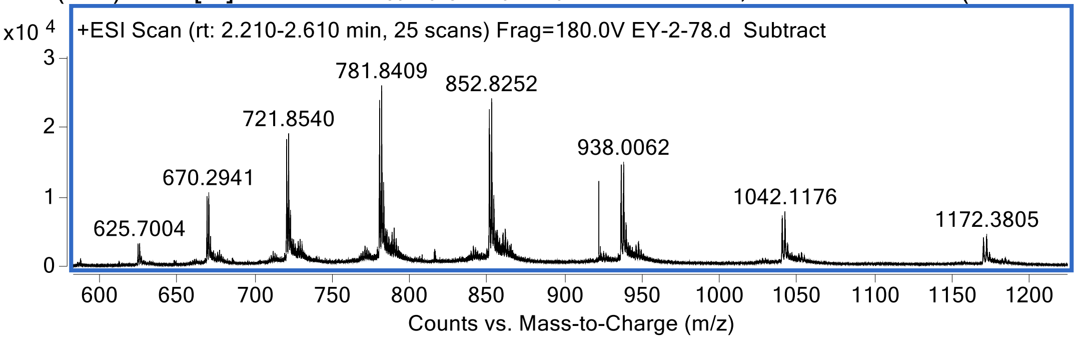


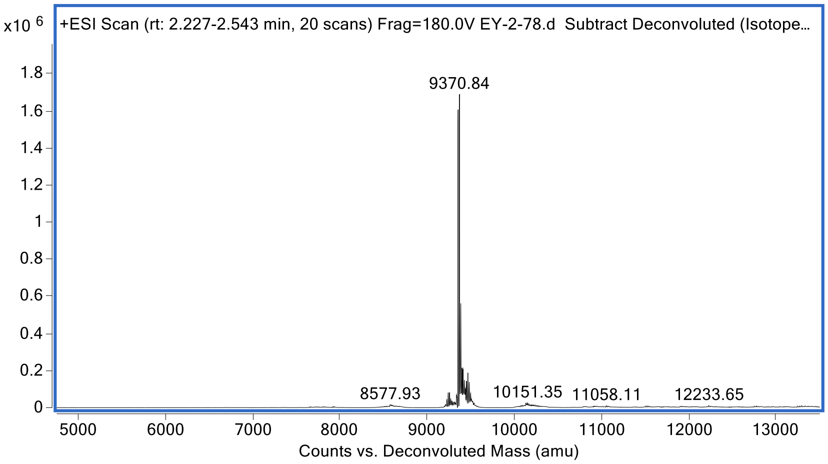

*Cy5-mISG15$_{CTD}$[R153Agb]-PA*

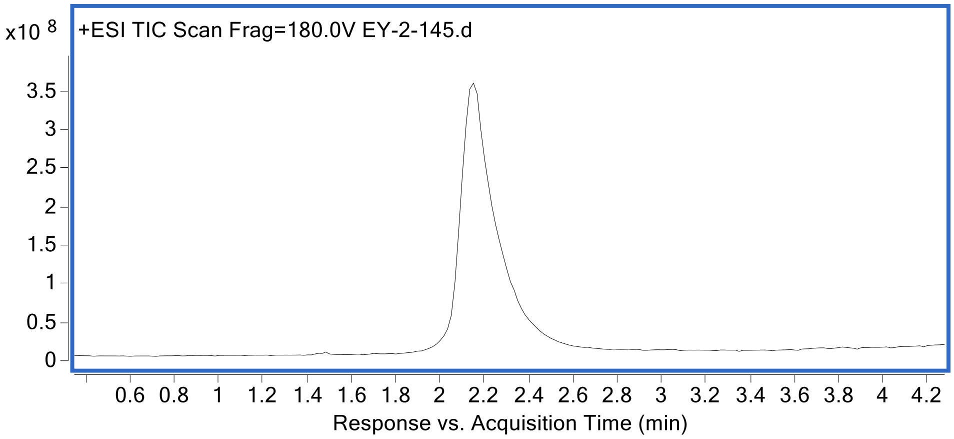


HRMS (ESI): m/z [M] calc. for $C_{429}H_{671}N_{116}O_{116}S$: 9341.8425, found 9341.24.

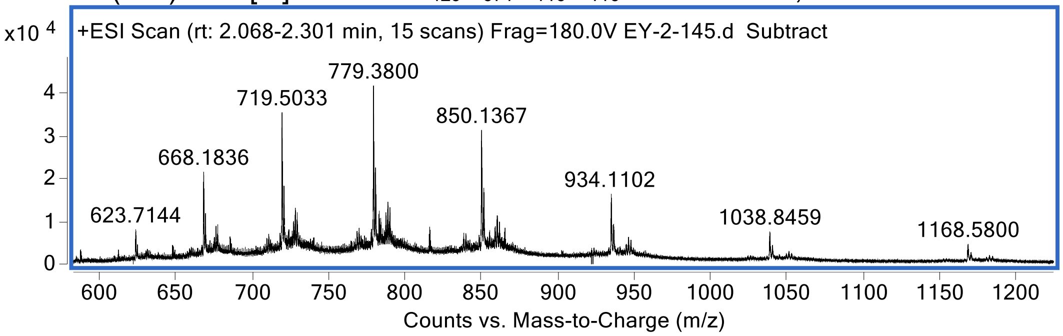


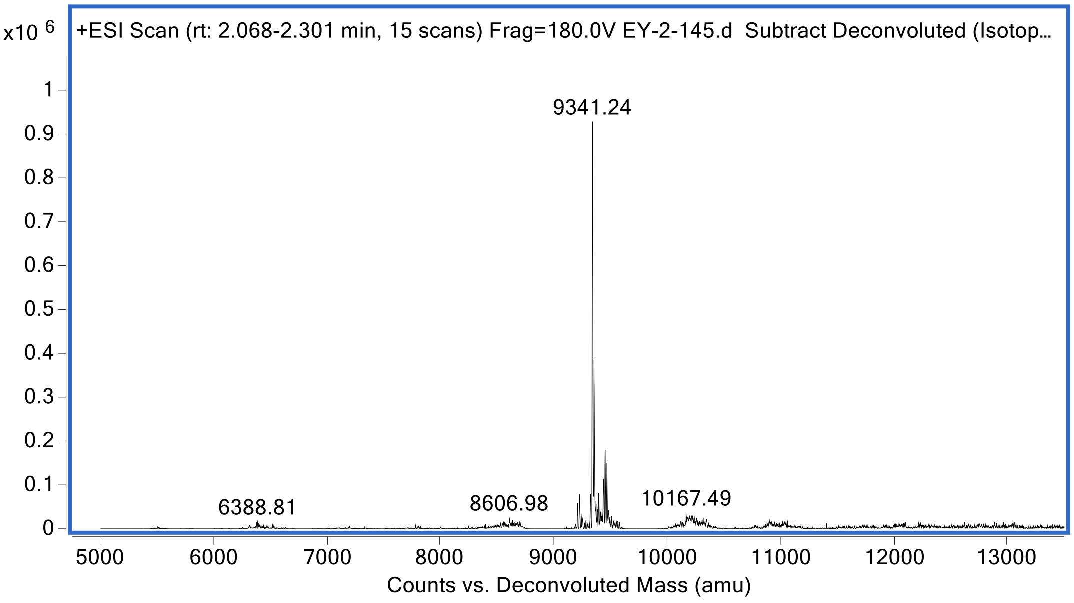

*Cy5-mISG15$_{CTD}$[R153F(guan)]-PA*

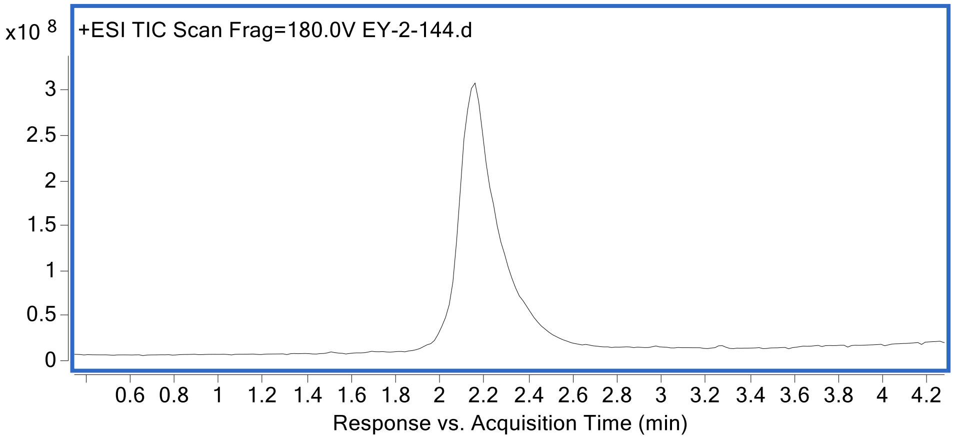


HRMS (ESI): m/z [M] calc. for $C_{434}H_{673}N_{116}O_{116}S$: 9403.9135, found 9403.33.

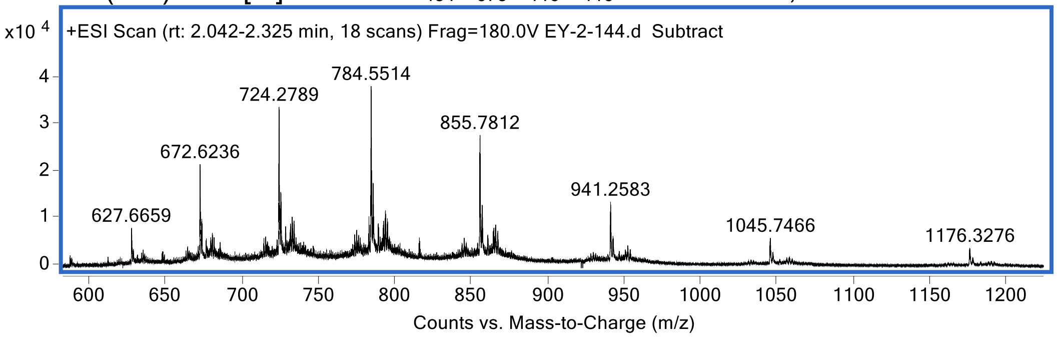


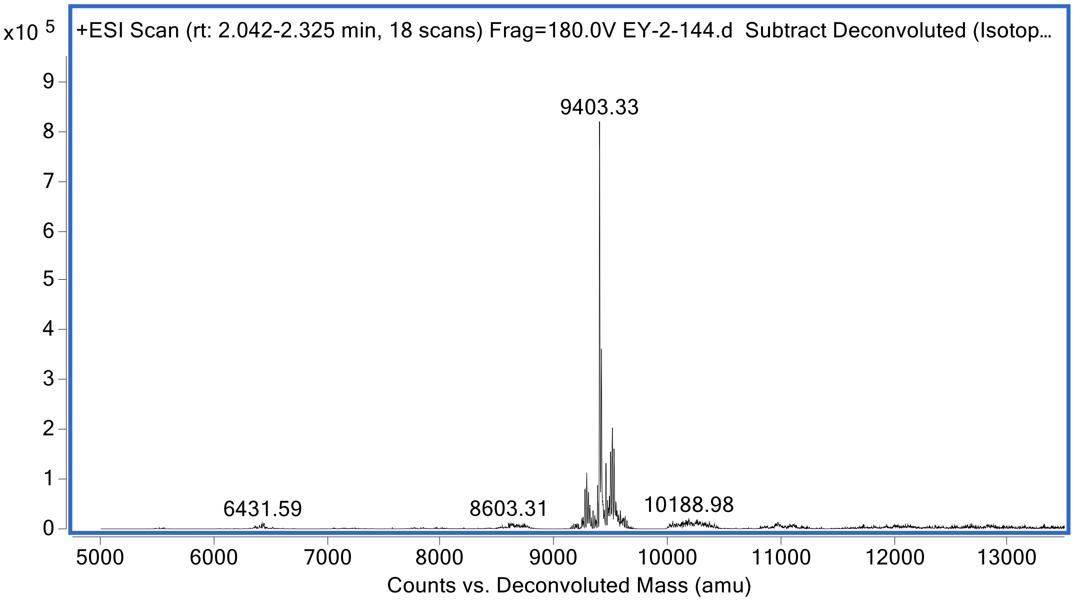

*Cy5-mISG15$_{CTD}$[H90F]-PA*

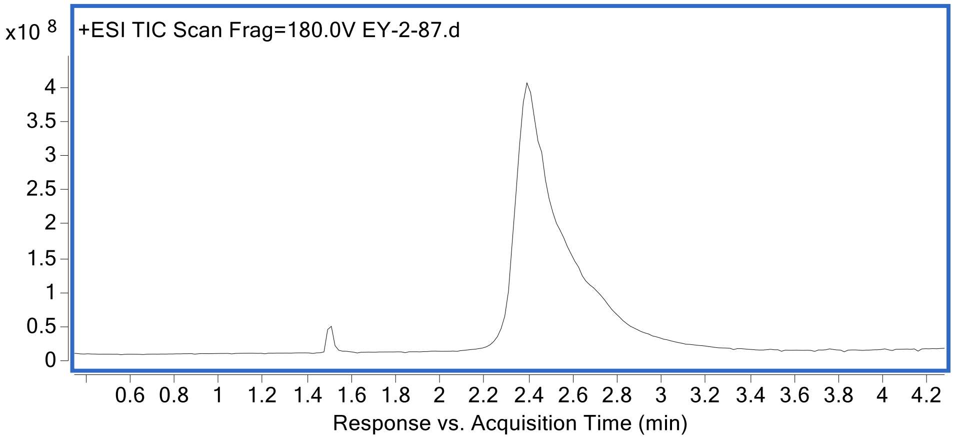


HRMS (ESI): m/z [M] calc. for $C_{433}H_{675}N_{114}O_{116}S$: 9365.9045, found 9381.35. (+15.4455)

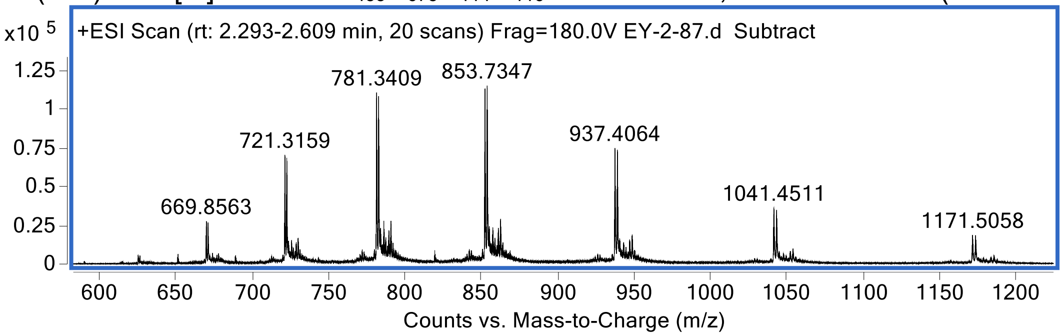


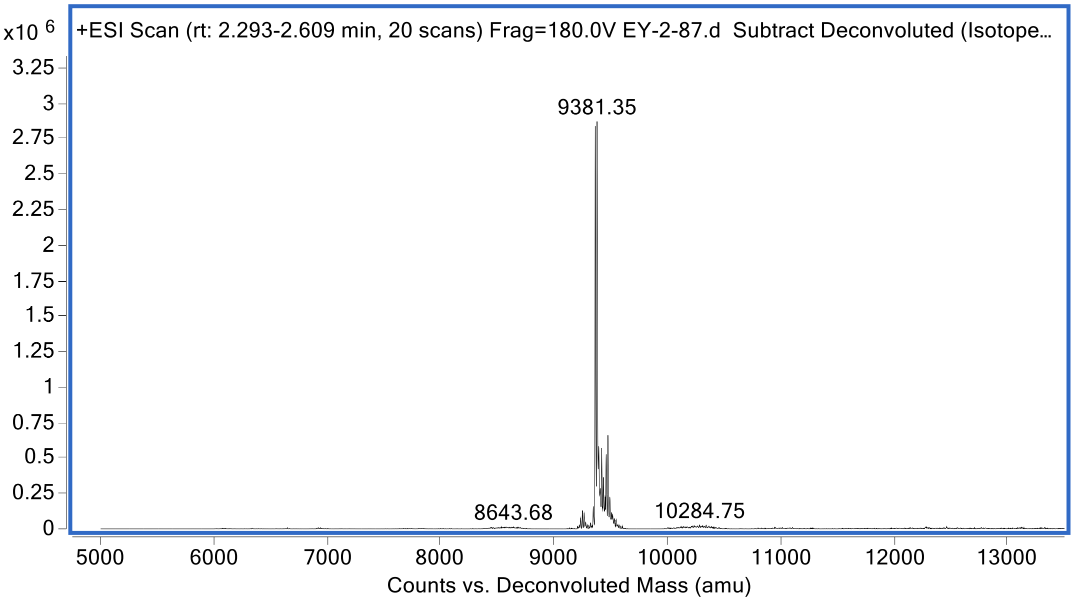

*Biotin-mISG15$_{CTD}$[WT]-PA*

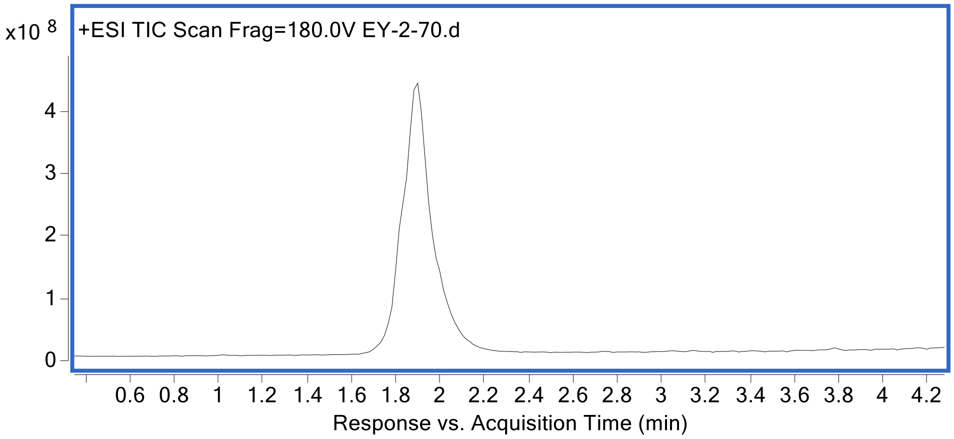


HRMS (ESI): m/z [M] calc. for $C_{414}H_{661}N_{117}O_{118}S_2$: 9229.6630, found 9246.12 (+16.457)

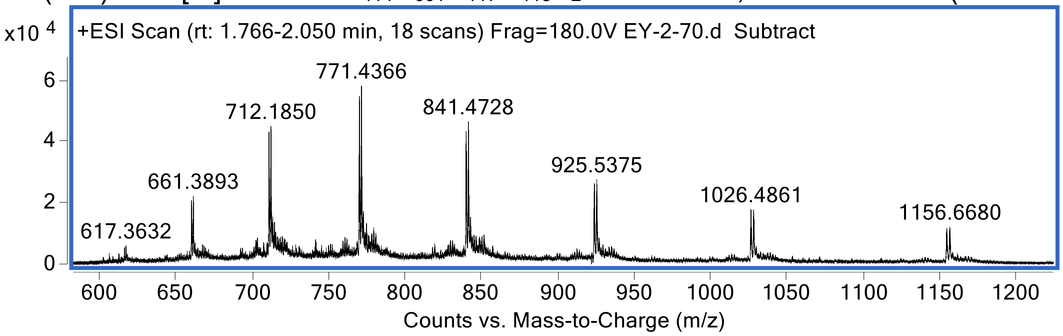


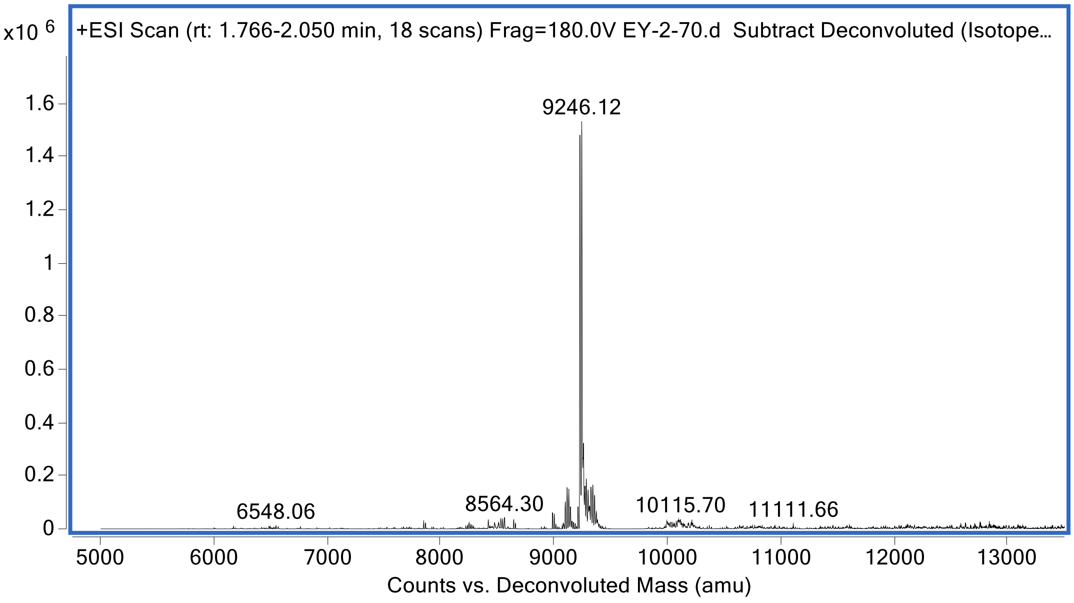

*Biotin-mISG15$_{CTD}$[R153Agb]-PA*

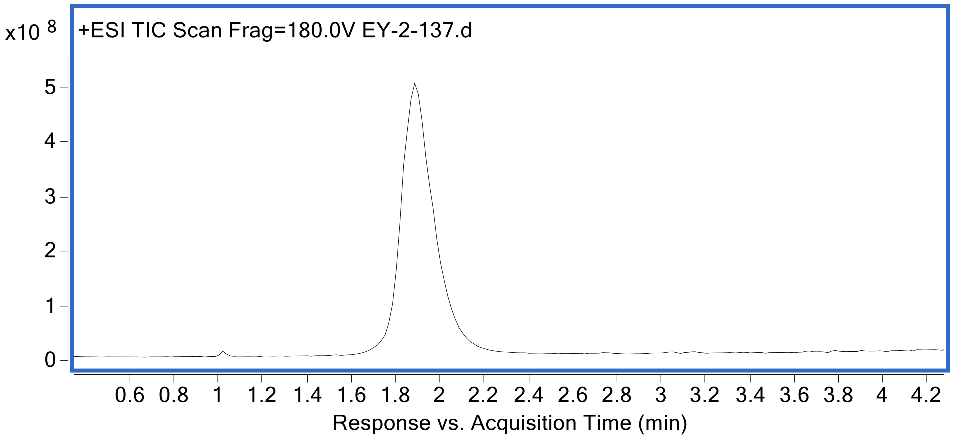


HRMS (ESI): m/z [M] calc. for $C_{413}H_{659}N_{117}O_{118}S_2$: 9215.6360, found 9216.15.

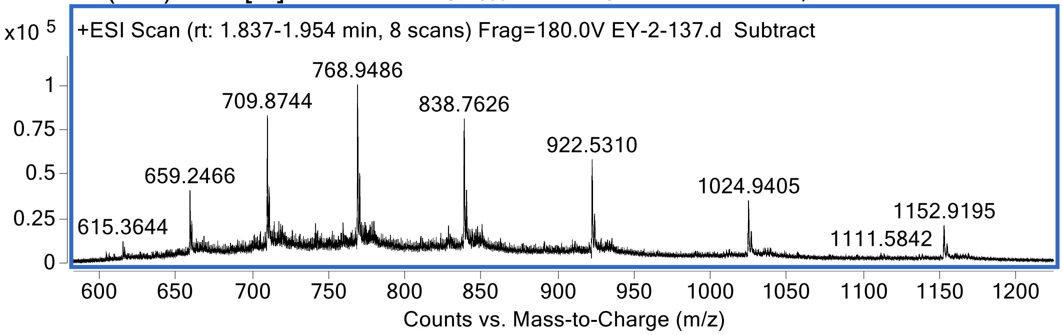


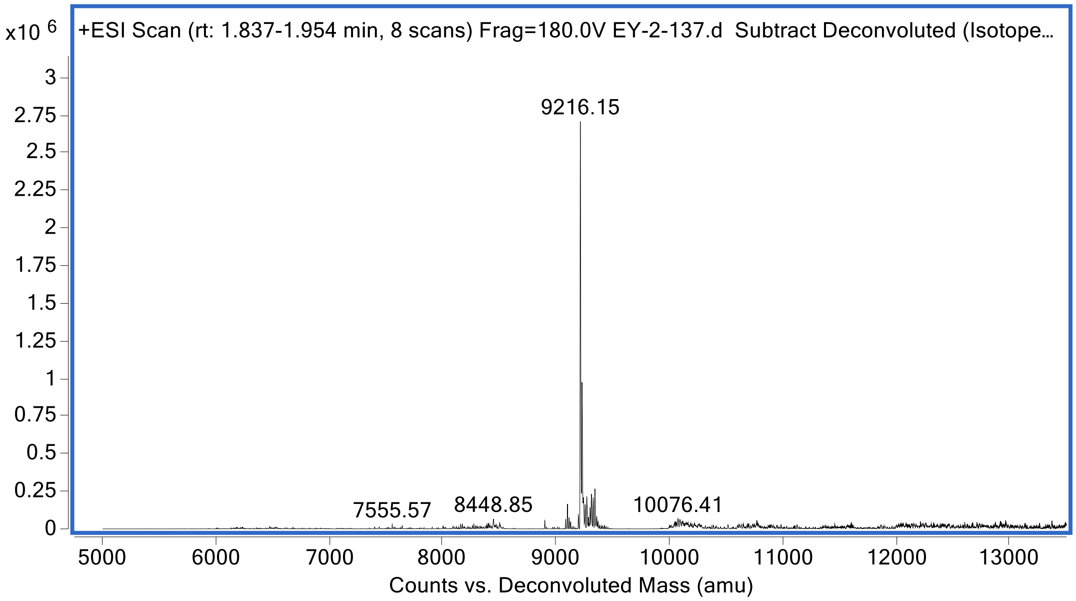

*Biotin-mISG15$_{CTD}$[R153F(guan)]-PA*

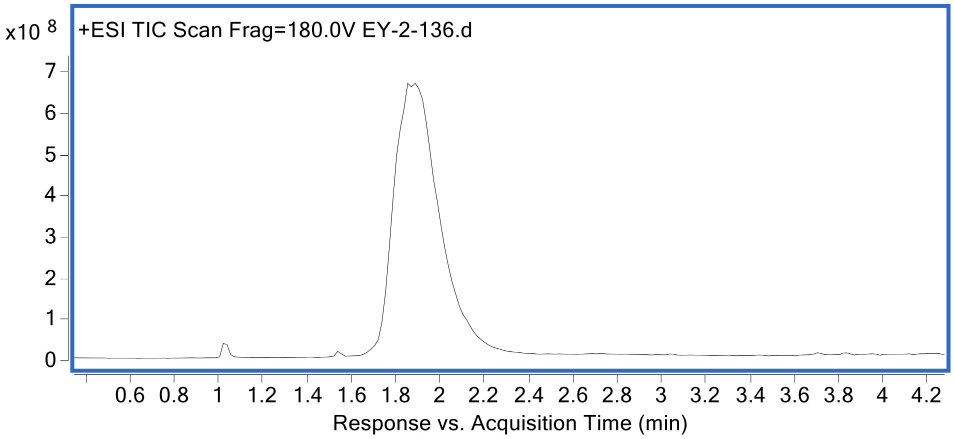


HRMS (ESI): m/z [M] calc. for $C_{418}H_{661}N_{117}O_{118}S_2$: 9277.7070, found 9278.21.

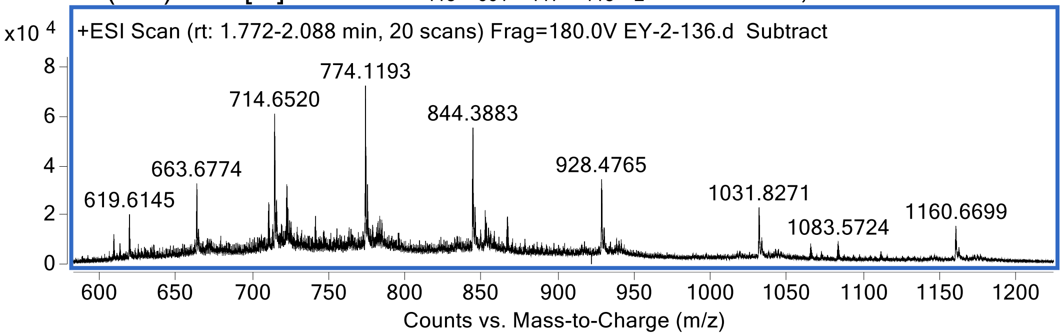


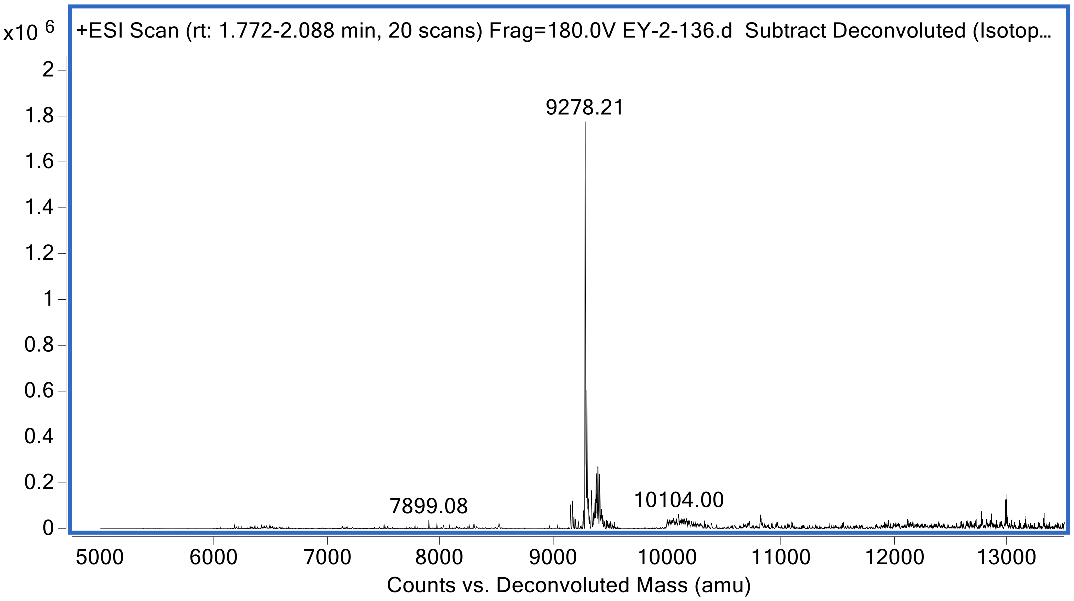
